# Commensal taxa in gut microbiota limit antibiotic resistance during extended oral antibiotic use

**DOI:** 10.1101/2025.08.13.670183

**Authors:** Erika L. Cyphert, Chongshan Liu, Victoria T. Chu, Aditi Dubey, Minying Liu, Zhe Zhong, James R. Cockey, Eva C. González Díaz, Angie L. Morales, Jacob C. Nixon, Matthew Garcia, Susana Zeng, Sidhant Rohatgi, Joan Wong, Ritwicq Arjyal, Honey Mekonen, Norma Neff, Jennifer Lee, M. Kyla Shea, Xueyan Fu, Sarah L. Booth, Cynthia A. Leifer, Ankur Singh, Charles R. Langelier, Christopher J. Hernandez

## Abstract

Certain bacterial infections, such as those involving prosthetics, can require antimicrobial therapy over months to years, potentially increasing the burden of antimicrobial resistance. Here we longitudinally track the antimicrobial resistome in mice during continuous antibiotic dosing over 21 months. The burden of antibiotic resistance genes (ARGs) initially increases, but, surprisingly, declines in later months, approaching levels observed in untreated animals. ARG burden is regulated by taxonomy and declines as ARG-harboring taxa that initially bloom are replaced by commensals. Furthermore, we find that the dynamics of antibiotic-induced ARG burden are influenced by age-related differences in microbial taxonomy and can be removed by fecal microbiota transplantation. We show that commensals may regulate the resistome by limiting the growth of ARG-harboring taxa, thereby providing antimicrobial expansion resistance.

## Main Text

Antibiotic-resistant infections are a global threat responsible for an estimated 1.5 million deaths annually (*1*). Widespread antibiotic use exerts selective pressure that drives the emergence of resistance among both commensal and pathogenic bacterial species (*2*). Collectively, antimicrobial resistance genes (ARGs) in an ecosystem are known as the “resistome.” While most antibiotic resistance mechanisms are naturally present, antibiotic use in humans and domestic animals has increased the collective reservoir of antibiotic resistance (*3–4*). Within hosts, the gut microbiome is the largest population of microbes and serves as an ecosystem for ARGs (*5–8*).

The abundance and richness of ARGs in the gut microbiome increases immediately after the start of oral antibiotics (*8–11*). A short course of oral antibiotics (e.g. < 10 days) results in changes in the composition of the microbiome and ARG abundance that reverses within several weeks to months after cessation (*8, 12–14*). However, there are certain clinical conditions in which prolonged antibiotic therapy of months to years is required, such as infections of prosthetic materials, endocarditis, or mycobacterial infections including tuberculosis (*15–18*). Although prolonged antibiotic exposure during infection treatment is presumed to select for accumulation of resistance, this has not been rigorously studied with respect to the microbiome resistome (*18, 19*).

Here, we use a mouse model to longitudinally assess community dynamics within the gut microbiome over 21 months of exposure to oral antibiotics. Surprisingly, after a year of continuous antibiotic dosing, the burden of ARGs decreases toward levels seen in treatment-naïve mice primarily due to shifts in taxonomic abundance. Furthermore, the dynamics of ARG abundance depend on age at antibiotic initiation; the transient increase in ARG abundance is shorter when antibiotics are initiated late in life. Lastly, we show that the abundance and richness of ARGs can return to baseline through administration of a fecal microbiota transplant (FMT) from treatment-naïve mice. Collectively, our findings illustrate how members of a microbial community with intrinsic resistance can regulate the resistome, thereby providing antimicrobial expansion resistance in a community.

### Abundance of antimicrobial resistance genes declines during prolonged exposure to oral antibiotics

To capture the dynamics of the microbiota community in mice throughout life, fecal samples were collected from mice longitudinally, from one month of age until 22 months of age (near the natural lifespan), to observe changes in the gut microbiome during normal aging (Unaltered) or during sustained dosing with oral antibiotics in drinking water (the beta-lactam ampicillin and the aminoglycoside neomycin, Continuous Dose; Fig. 1A). Only a portion of the microbiota is susceptible to one or both antibiotics; hence, many taxa are not directly affected by the applied antibiotics, as is true with most antibiotics used clinically (*21*). Chronic administration of these antibiotics did not have detectable effects on gut permeability, serum biomarkers of inflammation or systemic immunity (fig. S1-2, table S1-2).

**Fig. 1.**
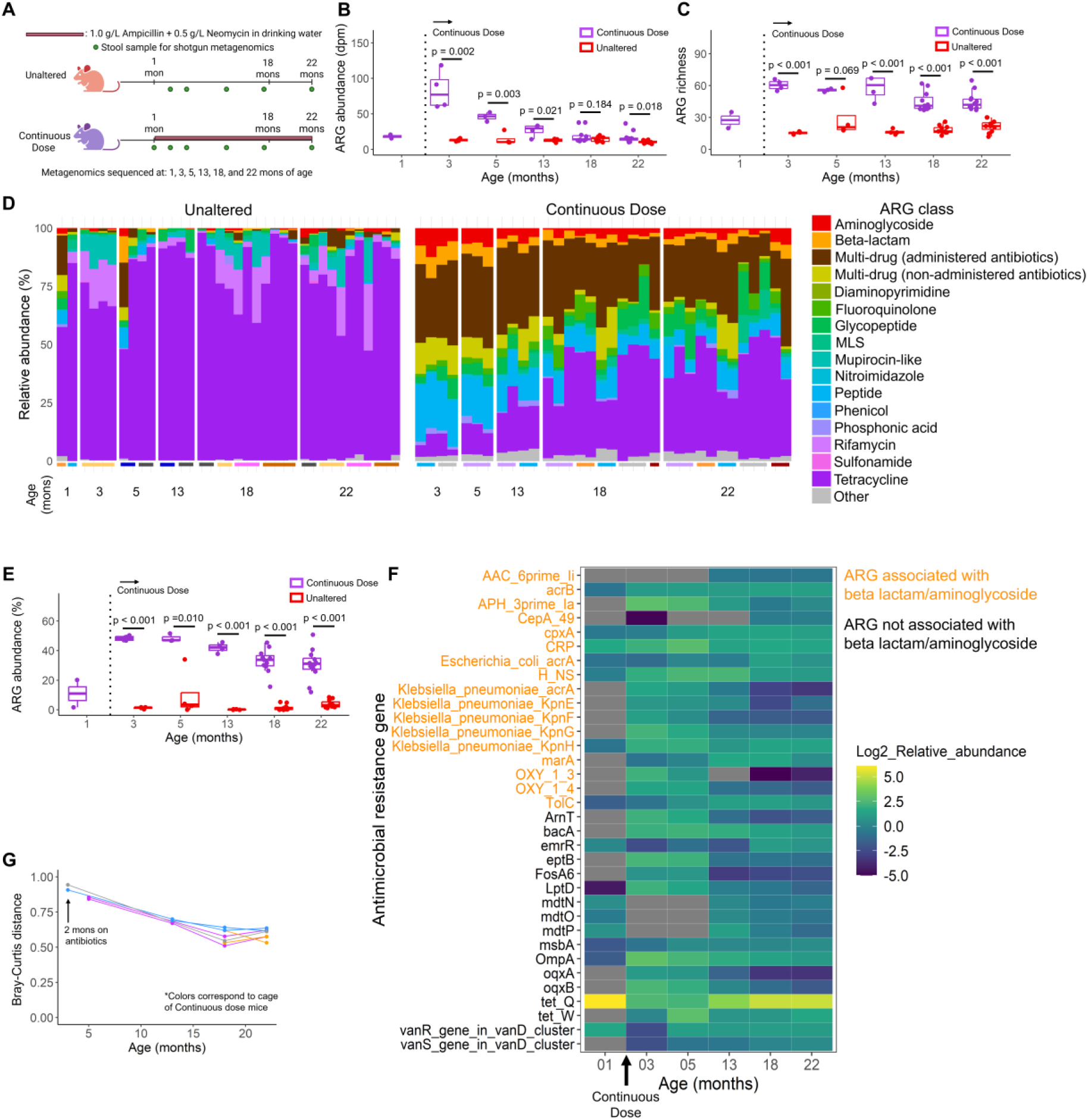
Abundance of antimicrobial resistance genes declines during prolonged exposure to oral antibiotics. (**A**) Study design. (B) Changes in ARG abundance over time are shown. (C) Changes in ARG richness are shown. (D) Bar charts for each animal show changes in ARGs. Colors at the bottom represent the same cage. (E) Two months after the start of antibiotic exposure < 50% of the ARG abundance conferred resistance to beta-lactams or aminoglycosides. (F) The increase in ARG richness was primarily due to the sustained presence of ARGs that were first detected shortly after the start of antibiotic dosing. (G) Longitudinal tracking in individual mice showed changes in the Bray-Curtis resistome composition over time that decreased. Lines connect data points from the same individual and represent Bray-Curtis distance to baseline (1 month of age prior to antibiotic exposure); colors correspond to the cages as shown in D. Results from female mice are shown here, the corresponding results from male mice are in Figure S3. Whiskers of the box plots represent the range of the data and the boxes represent the middle 50% of data.

In the first few months of continuous antibiotic dosing, ARG abundance and diversity increased, peaking at three months of age. The abundance of ARGs subsequently declined, approaching levels similar to those of the Unaltered group by 18 months of age (Fig. 1B, fig. S3). The richness of ARGs also peaked at three months of age and declined thereafter, but remained elevated relative to the Unaltered group for the remainder of the experiment (Fig. 1C, fig. S3).

The enrichment of ARGs only partially reflected the administered antibiotics (table S3-4). Prior to antibiotic exposure (at one month of age), the abundance of ARGs was low and primarily consisted of tetracycline resistance (*tet(Q)* > 90% relative abundance); ARGs for beta-lactam, aminoglycoside and multidrug resistance genes together were less than 4% of relative abundance. Two months after the start of antibiotic exposure, ARG abundance was increased, primarily due to enrichment of ARGs conferring resistance to the administered antibiotics (49.29 ± 2.18% associated with beta-lactam, aminoglycoside or multidrug resistance genes effective against either drug). By 22 months of age, only 26.28 ± 9.75% of ARG abundance was associated with applied antibiotics (as compared to 3.18 ± 2.27% in Unaltered mice) (Fig. 1D-E, fig. S3-4). These findings show that after years of continuous exposure to antibiotics, the majority of ARGs in a gut microbial community do not confer resistance to the applied antibiotics.

Although the abundance of ARGs decreased late in life in the Continuous Dose group, ARG richness remained elevated. The increase in ARG richness was due to the presence of ARGs that were not detected prior to the start of dosing (Fig. 1F, fig. S3). Several ARGs were first detected months after the start of antibiotic exposure. These “new” ARGs conferred resistance to multiple antibiotic classes including beta-lactam, aminoglycoside, tetracycline, and fluoroquinolone (Fig. 1, fig. S3). Additionally, during continuous dosing, the resistome composition shifted over time to be more similar to baseline as indicated by reductions in Bray-Curtis distance (Fig. 1G, fig. S3). Together, these findings show that during prolonged exposure to antibiotics, the majority of enriched ARGs confer resistance to unrelated antibiotic classes and total ARG abundance can be similar to that in Untreated individuals.

### Taxonomic changes explain declines in ARG abundance during prolonged oral antibiotics

The observed ARGs were attributed to several different taxa based on the CARD ontology, leading us to examine the changes in taxonomic composition in more detail. Consistent with prior work (*21, 22*), oral antibiotics resulted in a drastic change in gut microbial composition compared to mice without antibiotic exposure. Although antibiotic perturbation was continuous, the gut microbial population was dynamic, with significant differences observed in the taxonomic composition between three and 22 months of age (PERMANOVA: female – p < 0.001, male – p = 0.003; Fig. 2A, fig. S5). Specifically, the Bray-Curtis distance of taxonomic composition in individual animals decreased over time, becoming more similar to the Unaltered baseline (Fig. 2B, fig. S5).

**Fig. 2.**
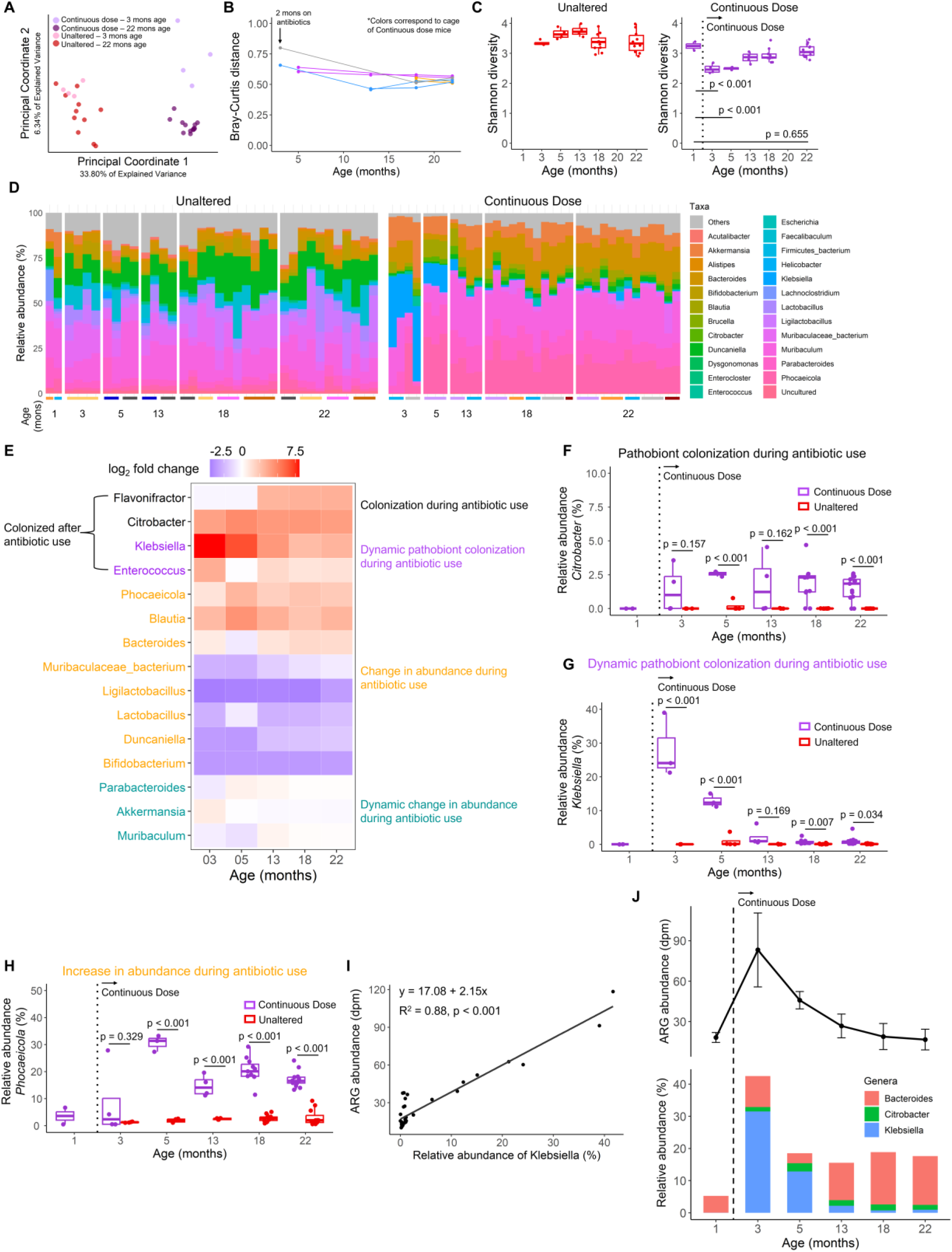
Taxonomic changes explain declines in ARG abundance during prolonged oral antibiotics. **(A)** Continuous oral antibiotics resulted in a rapid change in gut microbial composition compared to Unaltered mice. (B) During continuous dosing, the microbial composition shifted slightly toward the Unaltered state seen in the Bray-Curtis distance from baseline. (C) Shannon diversity initially decreased in Continuous Dose mice but recovered afterwards. (D) Genus-level analysis revealed complex dynamics of taxa during antibiotic dosing. (E) Log-fold change of most abundant genera during antibiotic dosing relative to one month of age before antibiotics. Representative pathobionts that (F) colonized, (G) dynamically colonized; or (H) increased in abundance during antibiotic use. (I) The relative abundance of *Klebsiella* is strongly correlated with ARG abundance in Unaltered and Continuous Dose mice (Pearson correlation: 0.88). (J) Shifts in the relative abundance of ARG harboring taxa (*Bacteroides*, *Citrobacter*, *Klebsiella*) directly align with shifts in ARG abundance. Whiskers in the line plot represent the standard deviation of data. Whiskers of the box plots represent the range of the data and the boxes represent the middle 50% of data. Results from female mice are shown here; corresponding results from male mice are in Figure S5.

Short term antibiotic dosing reduces the alpha diversity of the gut microbiome (*22*). Consistent with this fact, in the two months after the start of continuous antibiotic dosing, Shannon diversity was reduced relative to baseline (Fig. 2C). After 12 months of continuous dosing, however, Shannon diversity had unexpectedly increased in females, nearly reaching baseline by 22 months of age (Fig. 2C, in males, Shannon diversity remained low, fig. S5). The increase in Shannon diversity in females was attributed to the increased abundance of more than 100 genera. We speculate that Shannon diversity recovered in females because females tend to have a greater microbiome diversity than males (*23*). Genus-level analysis revealed complex dynamics of taxonomic composition during continuous antibiotic dosing (Fig. 2D-E, fig. S5-6). The dynamic changes in taxonomic abundance followed four different responses to antibiotic exposure: colonization, dynamic colonization, sustained change in abundance, and dynamic change in abundance. Representative genera that followed these trends include *Citrobacter*, *Klebsiella*, and *Phocaeicola* (Fig. 2F-H, fig. S5).

Prior to antibiotic exposure, *Muribaculaceae spp.* occupied the largest proportion of taxonomic abundance (29.81 ± 2.13%). Shortly after the onset of antibiotic exposure, the abundance of *Muribaculaceae spp.* declined (14.27 ± 8.86%). Concurrently, the relative abundance of *Enterobacteriaceae* rapidly increased and peaked at three months of age (29.35 ± 8.67%) and later declined to 2.66% by 22 months of age (still greater than the pre-antibiotic baseline of 0.26 ± 0.32%). The changes in *Enterobacteriaceae* abundance in the Continuous Dose group were predominantly driven by two classic pathobionts that frequently harbor ARGs: *Klebsiella spp.* and *Citrobacter spp.* (table S5). The decline in *Enterobacteriaceae* coincided with the gradual increase in the relative abundance of *Bacteroides*. Before exposure to antibiotics, *Bacteroides* had a relatively low abundance (6.56 ± 3.16%). Prolonged exposure to antibiotics gradually increased the relative abundance of *Bacteroides* which peaked at 16.24 ± 2.97% in females. *Bacteroides* are not susceptible to aminoglycosides and commonly carry resistance against beta-lactams and aminoglycosides (*24, 25*).

The changes in taxa over time in the Continuous Dose group were substantially larger than the age-related changes observed in the Unaltered group, which experienced subtle changes in taxonomy during aging (table S6). The changes in taxonomic composition explained the majority of changes in ARG abundance. Changes in the relative abundance of *Klebsiella* alone explain more than 85% of the variance in total ARG abundance (Fig. 2I, fig. S5,7-8). Overall, the transient increase in ARG burden aligned with the combined relative abundance of *Klebsiella*, *Citrobacter*, and *Bacteroides* (Fig. 2J, fig. S5). These three genera harbor 82% of the differentially abundant ARGs (based on CARD ontology/pathogen of origin prediction). As the abundance of *Klebsiella* and *Citrobacter* increased upon exposure to antibiotics, the abundance of ARGs increased. When the abundance of *Klebsiella* decreased, ARG abundance also decreased but was partially replaced by ARGs associated with *Bacteroides*. Together, these results show that fluctuations in the abundance of ARGs during continuous antibiotic exposure are explained primarily by shifts in taxonomy as ARG-harboring organisms are replaced by commensals of varying susceptibilities.

### Age-related change in the gut microbiota influences ARG dynamics following initiation of oral antibiotics

The response of the gut microbiota to oral antibiotics is influenced by host age (*26*). As the composition of the gut microbiota is more malleable in younger than in older mice (*27, 28*), we hypothesized that oral antibiotics would have less effect on the microbiota and the resistome in older mice. To test this idea, we initiated oral antibiotics in mice who were treatment-naïve until 18 months of age (considered elderly in mice) (Fig. 3A).

**Fig. 3.**
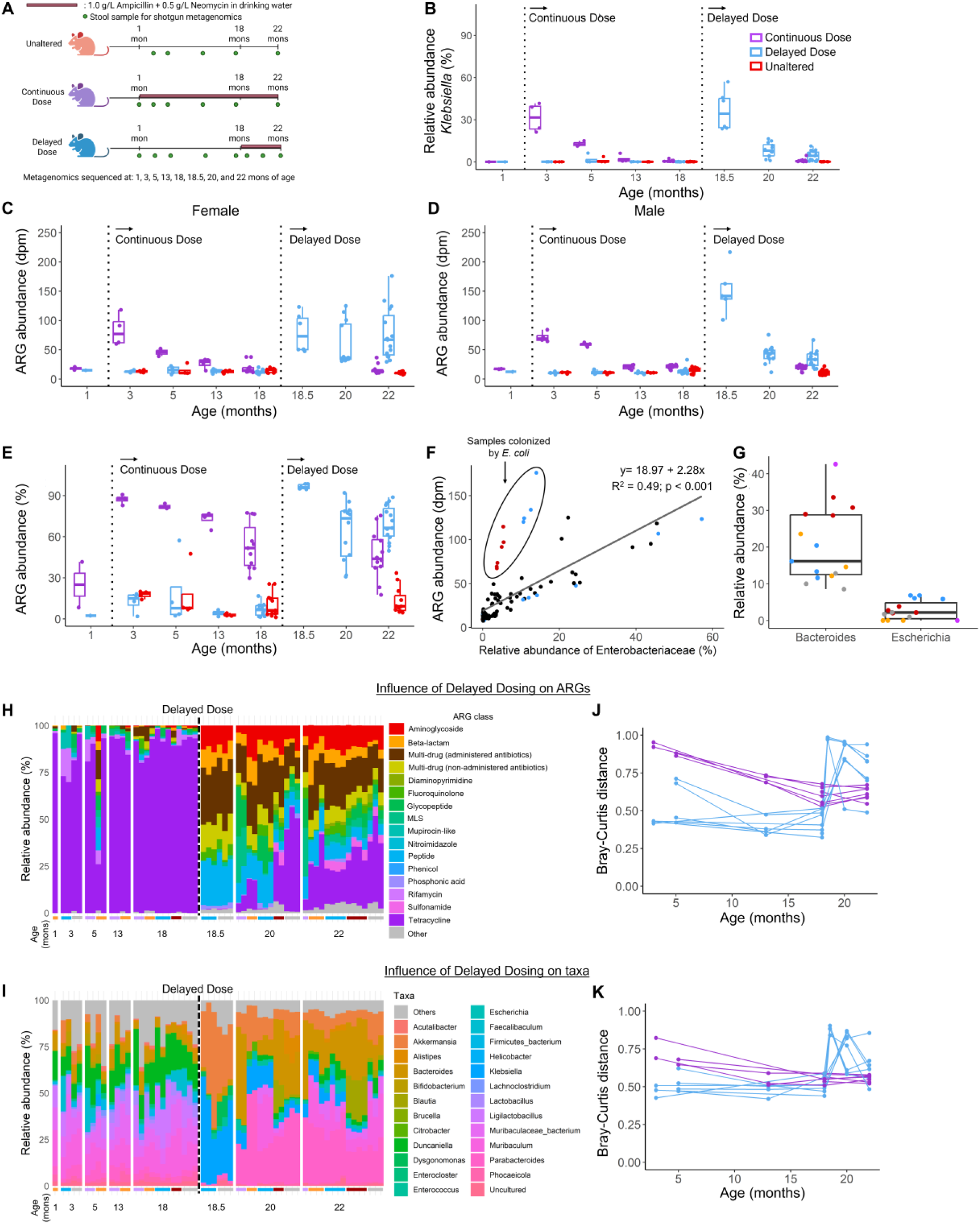
Age-related change in the gut microbiota influences ARG dynamics following initiation of oral antibiotics. **(A)** Study design. (B) Mice initiating antibiotic dosing at 18 months of age (Delayed Dose group) had a transient increase in *Klebsiella* and a more rapid decline in *Klebsiella* relative to younger mice in the Continuous Dose group that were exposed to the antibiotics for the same period. (C) In contrast to Continuous Dose mice, female Delayed Dose mice did not exhibit a decreased ARG abundance with prolonged antibiotic exposure. (D-E) Changes in ARG abundance in Delayed Dose mice. (F) The relative abundance of *Enterobacteriaceae* was strongly correlated with the abundance of ARGs with the exception of animals colonized with *E. coli*. (G) The relative abundance of *Bacteroides* and *Escherichia* in Delayed Dose mice (indicated by color). (H-I) The variability in the resistome and microbiota composition. (J-K) Variability in the Bray-Curtis distance from baseline (pre-antibiotic) in the Delayed Dose mice at 22 months of age was greater than that in Continuous Dose. Whiskers of the box plots represent the range of the data and the boxes represent the middle 50% of data. Additional results are shown in Figure S9.

The initiation of antibiotic dosing at 18 months of age (Delayed Dose groups) resulted in an increase in ARG abundance that was more heterogeneous than in Continuous Dose mice exposed to the antibiotics for the same period of time (Fig. 3C,D, table S8-9). Prior to antibiotic exposure, most ARGs were associated with tetracycline but after antibiotic exposure ARGs were primarily multidrug resistant and associated with aminoglycoside and beta-lactam (Fig. 3H, table S7). Unlike the Continuous Dose mice, however, the resistome of Delayed Dose mice showed more variability among animals within the same groups, as indicated by Bray-Curtis distance from baseline (Fig. 3J, fig. S9).

The initial shifts in taxonomy after the start of oral antibiotics followed a similar pattern to that seen in the Continuous Dose animals (Fig. 3I, fig. S9). As observed in the Continuous Dose groups, antibiotic dosing initiated a transient increase in the relative abundance of *Klebsiella*: The relative abundance of *Klebsiella* prior to antibiotic dosing was initially less than 1%, increased substantially within the first two months of antibiotic exposure, then declined (Fig. 3B, fig. S9). The decline in *Klebsiella* abundance in Delayed Dose mice followed an exponential decay that was much faster than in Continuous Dose mice exposed to antibiotics for the same duration (decay rate constant between two and four months of dosing 0.537/month in Delayed Dose mice versus 0.184/month in Continuous Dose, p < 0.001) (Fig. 3B) suggesting a more resilient microbiome in the aged Delayed Dose mice, consistent with prior reports and clinical findings (*8*, *29*). The taxonomic beta diversity in the Delayed Dose mice after the start of antibiotics was more variable, mirroring the differences in ARG diversity (Fig. 3J-K, fig. S9).

The relative abundance of *Klebsiella* was strongly correlated with ARG burden in the Delayed Dose mice but there were several outliers that showed larger than expected ARG burden (Fig. 3F). The outliers were a subset of animals with increased abundance of the ARG carrier *E. coli* and ARGs associated with *E. coli* (Fig. 3G, fig. S10). The presence of ARGs harbored by *E. coli* contributed to the maintenance of a higher total ARG burden in Delayed Dose mice, even when *Klebsiella* abundance declined (fig. S11). These findings further show that taxonomic differences drive variations in ARG abundance, even in aged hosts.

A secondary cause for variability in ARG abundance in Delayed Dose mice was the accumulation of ARGs in the microbiota in the months prior to the start of antibiotic dosing. At one month of age, more than 95% of ARG abundance was associated with tetracycline. By 18 months of age, the abundance of ARGs associated with tetracycline declined; *tet(Q)*, accounted for more than 70% of ARG abundance at one month of age but only 47% of ARG abundance at 18 months (fig. S12). Several “new” ARGs not detected at one month of age, but present at 18 months of age confer resistance to aminoglycoside, beta-lactam, glycopeptide, MLS, mupirocin-like, peptide, and multidrug resistance (Fig. 3H, fig. S10). These “new” ARGs were likely associated with changes in taxonomy (fig. S13). This observation is consistent with recent cross-sectional studies in aging humans that show increases in ARG abundance in older individuals (*30*). Although increased abundance of ARGs in older humans is often attributed to the accumulated effects of doses of antibiotics throughout life (*22*), our findings in mice suggest such shifts can occur in the absence of antibiotic exposure. Collectively, our findings show that subtle changes in the gut microbiota during aging can change the response to oral antibiotics in terms of taxonomic composition and associated ARGs.

### Fecal microbiota transplantation mitigates collateral ARGs acquired through lifelong antibiotic exposure

The accumulation of ARGs in response to antibiotic exposure has the potential to provide a reservoir of antibiotic resistance that can be transmitted through communities or complicate management of bacterial infections (*31*). Fecal microbiota transplantation (FMT) has been proposed as a means of displacing resistant organisms from the microbiome (*9*). We thus evaluated the utility of FMT for reducing ARG burden in mice continuously exposed to antibiotics from one to 18 months of age. At 18 months of age, antibiotic dosing was suspended, and mice received FMT from age- and sex-matched donors with an Unaltered microbiome (Initial Dose group, Fig. 4A). Within two weeks after the FMT, the taxonomic composition of the gut microbiota became indistinguishable from that of similarly aged treatment-naïve animals (Fig. 4B, fig. S14-15). Similarly, the composition, abundance, and richness of ARGs matched that of age- and sex-matched Unaltered animals (Fig. 4C-E, fig. S14). After the FMT, species that had colonized as a result of antibiotic dosing were no longer detectable after FMT (Fig. 4G, fig. S14). Therefore, the introduction of treatment-naïve microbiota not only replenished the microbial taxa to an Unaltered state but also removed ARGs that were enriched during prolonged antibiotic dosing.

**Fig. 4.**
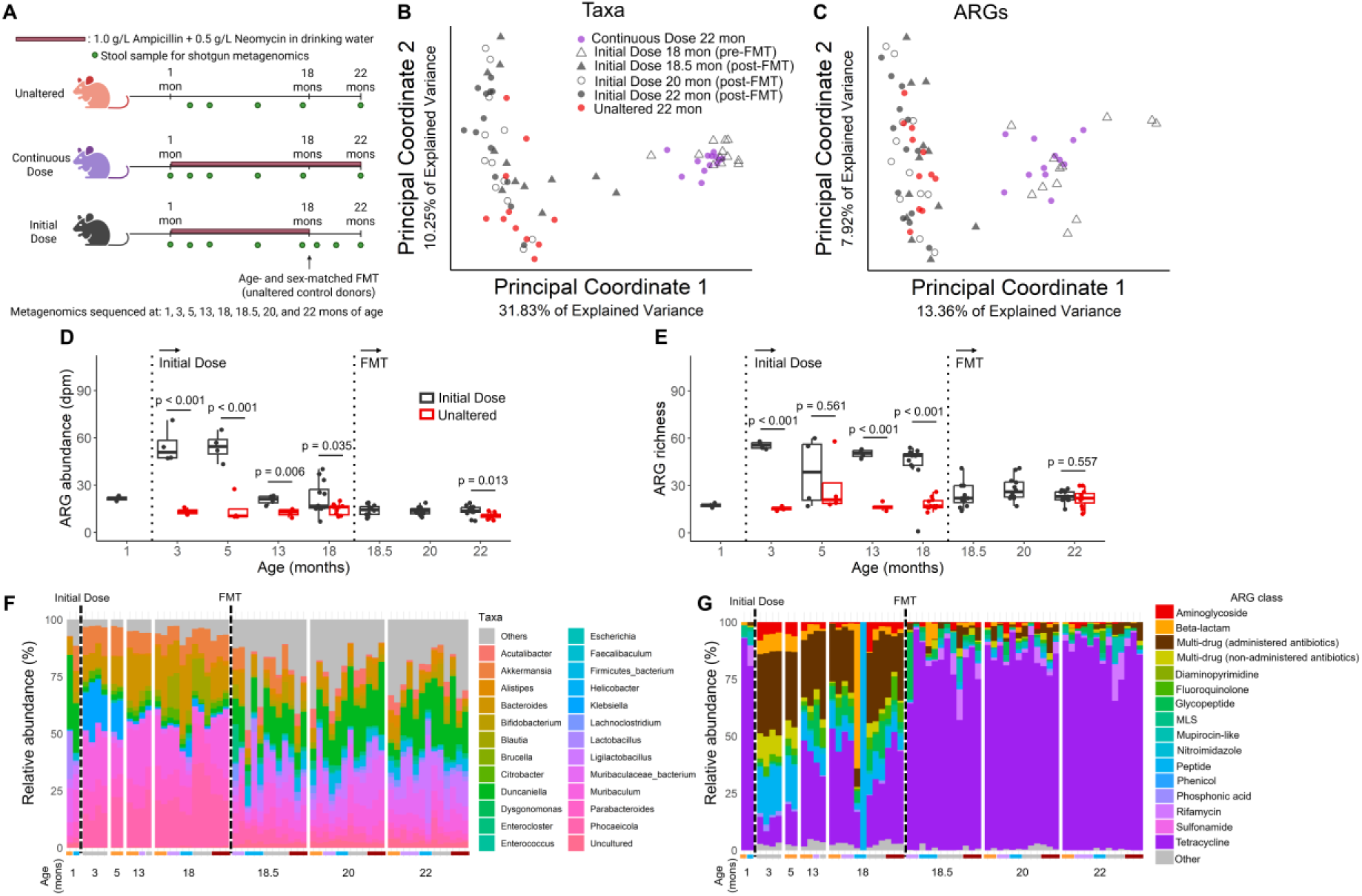
Fecal microbiota transplantation mitigates collateral ARGs acquired through lifelong antibiotic exposure. (**A**) Study design. (B-C) Receiving the FMT from the Unaltered donor at 18 months of age restored the microbiota and resistome composition to its Unaltered state. (D-E) The total abundance and richness of ARGs decreased, returning to the level of Unaltered mice after receiving the FMT. (F-G) Composition of the microbiota and resistome prior to and after receiving the FMT. Whiskers of the box plots represent the range of the data and the boxes represent the middle 50% of data.

## Discussion

Extended exposure to antibiotics has been implicated in promoting antibiotic resistance by selecting for resistant organisms and thus increasing the abundance of ARG in the microbiome. Antibiotic stewardship strategies that limit excessive or inappropriate antibiotic use reduce the incidence of antibiotic resistant infections (*32*). Certain types of infections, however, require prolonged antibiotic treatment for months or years (*15–18*), raising the risk that commensals and/or pathogens in the gut microbiota will develop antibiotic resistance. Our findings demonstrate that the ARG burden increases most strikingly following initiation of antibiotic treatment, and then recedes after months of exposure, but never fully normalizes. This likely represents an initial fitness advantage of taxa harboring relevant ARGs that ultimately get outcompeted by community members with intrinsic resistance or other growth advantages in the presence of antibiotics. Our findings illustrate how members of the community that are not susceptible to an antibiotic can provide expansion resistance limiting the abundance of ARGs and emergence of resistant pathogens.

Studies of antibiotic resistance in the gut microbiota report increases in ARG abundance and richness during and immediately after short-term (fewer than 10 days) antibiotic dosing (*22*, *33–35*) and/or cross-sectional studies in humans (*36–38*). These studies have shown that short-term antibiotic exposures can have a lasting effect on the gut microbiota and resistome, although some components of the microbiota and resistome eventually return to pre-treatment states (*8*, *22*, *39–42*). Consistent with prior reports (*10*, *43*), we observed increases in ARG abundance and reductions in taxonomic diversity soon after antibiotic dosing. However, unlike prior reports that have been cross-sectional, our study is unique in performing a longitudinal analysis for most of the natural lifespan of the mice. We were therefore able to observe dynamic changes in the gut microbiota and ARG abundance over almost two years of dosing. Our findings show that the early increases in ARG abundance can eventually fade.

A common concern related to excessive antibiotic dosing is the possibility that horizontal gene transfer could confer resistance to antibiotics among organisms in the gut microbiota (*3*, *44*, *45*). Although we did not directly examine horizontal gene transfer, our finding that fluctuations in taxonomic abundance explain over 85% of ARG abundance suggests that the majority of the ARGs in our study were taxonomically stable, making it unlikely that horizontal gene transfer had a significant effect on the overall ARG profile.

Our findings that FMT eliminates the detection of ARGs and taxa acquired after prolonged antibiotic use aligns with clinical findings that FMT reduces the abundance of ARGs (*9*, *46*, *47*). Collectively, our findings support the idea that FMTs can reduce the abundance of ARGs even after prolonged antibiotic exposure. However, our findings highlight the importance of the microbiota before perturbation.

There are some important limitations to our study. First, identification, classification and association with host taxa of ARGs was limited to the databases used. Although the CZ-ID pipeline uses the most updated National Center for Biotechnology Information (NCBI) and Comprehensive Antibiotic Resistance Database (CARD) databases (*48*), it only includes ARGs that have been previously sequenced and classified and is therefore biased to the most commonly studied organisms (e.g., the *Enterobacteriaceae* family). It is unclear how much of our findings are specific to the murine host, diet, the breadth of the antibiotic spectrum, or the specific antibiotics used in this study.

Our findings show that the impact on ARG abundance is greatest in the first weeks after antibiotic exposure. As a result, risks of transmission of antibiotic-resistant taxa and/or antibiotic-resistant infections from the gut microbiota may be greatest in the first few weeks to months of exposure, while steadily declining if antibiotic dosing continues for longer periods. In situations requiring prolonged antibiotics, the risk presented by increased ARG abundance in the gut microbiota may decline after the first few months of dosing. Our findings demonstrate that the presence of commensal microbes that are not sensitive to the dosed antibiotic provides a limit to the accumulation of antibiotic resistance within the community, an effect we term expansion resistance. Taken together, our results demonstrate that a more diverse gut microbiome not only helps to promote colonization resistance but can also moderate the abundance of antibiotic-resistant microbes.

## Funding

National Institutes of Health grant R01AG067996 (CJH)

National Institutes of Health grant F32AG076244 (ELC)

National Institutes of Health grant 1R01AI181282-01A1 (AS)

USDA Agricultural Research Service Cooperative Agreement No. 58-8050-3-003 (ML, MKS, XF, SLB)

## Author contributions

Each author’s contribution(s) to the paper should be listed [we encourage you to follow the CRediT model]. Each CRediT role should have its own line, and there should not be any punctuation in the initials.

Examples:

Conceptualization: ELC, CL, VTC, CRL

Methodology: ELC, CL, VTC, AD, ML, ZZ, JRC, RA, HM, NN, JL, MKS, XF, SLB, CAL, AS, CRL

Investigation: ELC, CL, AD, ML, ZZ, JRC, ECGD, ALM, JCN, MG, SZ, JW, JL, MKS, XF, SLB, CAL, AS, CRL

Visualization: ELC, VTC, ML, ZZ, SR, CRL Funding acquisition: CJH

Project administration: CJH Supervision: CJH

Writing – original draft: ELC, VTC, CRL, CJH

Writing – review & editing: ELC, CL, VTC, AD, ML, ZZ, JRC, ECGD, ALM, JCN, MG, SZ, SR, JW, RA, HM, NN, JL, MKS, XF, SLB, CAL, AS, CRL, CJH

## Data and materials availability

Raw shotgun metagenomics fastq.gz files can be accessed on Dryad database.

**Supplementary Materials**

Materials and Methods

Figs. S1 to S15

Tables S1 to S10

## SUPPLEMENTARY

### Materials and Methods

#### Mouse strains

All animal procedures received prior approval from the local Institutional Animal Care and Use Committee. Male and female C57Bl/6J mice were purchased from Jackson Laboratory (Bar Harbor, ME, USA) and bred using trio breeding Pups were weaned at three weeks of age and housed by sex (n = 4/cage). Second-generation pups from breeders were used in the study. Sterile cages contained ¼-inch corn cob bedding (The Andersons’ Lab Beddin, Maumee, OH, USA), standard laboratory chow (Teklad LM-485 Mouse/Rat Sterilizable Diet Irradiated, Envigo Diets, Madison, WI, USA), reverse osmosis sterilized water *ad libitum*, and cardboard hut (Ketchum Manufacturing, Brockville, Canada).

#### Study design

Male and female pups were divided into four treatment groups (n > 15/sex/group) over two breeding rounds (same dams, two months apart). Treatment groups included: 1) Unaltered control from birth-22 months of age, 2) Continuous alteration from 1-22 months of age, 3) Initial alteration from 1-18 months of age that received an age- and sex-matched Unaltered donor fecal microbiota transplant at 18 months of age, and 4) Delayed alteration from 18-22 months of age only. Antibiotic dosing (microbiome alteration) consisted of reverse osmosis sterilized water or reverse osmosis sterilized water containing 0.5 g/L neomycin (N6386, Millipore Sigma, Burlington, MA, USA) and 1 g/L ampicillin (A0166, Millipore Sigma, Burlington, MA, USA) in light-sensitive water bottles. Fresh antibiotic water was replaced every three days over the duration of the study. Antibiotics were selected due to their poor oral bioavailability and limited spectrum of activity, capable of acting directly on gut microbes. Weekly bedding mixing was performed between cages of the same sex and treatment groups from 1-3 months of age. Specifically, a sample of dirty bedding was collected from each cage within a treatment group, pooled together and redistributed back to the cages to normalize gut microbiota composition across cages of the same treatment and sex (*49*).

#### FMT to antibiotic-treated mice

Male and female C57Bl/6J mice (n = ∼15/sex) that had been treated with ampicillin and neomycin in their drinking water from 1-18 months of age received a fecal microbiota transplant (FMT) at 18 months of age to repopulate microbes. Mice were fasted for three hours prior to receiving the transplant with *ad libitum* access to water. Fresh stool samples were collected from age- and sex-matched unaltered control mice with two pellets being placed in sterile 1.5 mL microcentrifuge tubes. Fecal samples were immediately transferred to an anaerobic chamber (Coy Laboratory Products, Grass Lake, MI, USA) and suspended in 1 mL anoxic PBS with 0.05% L-cysteine. Pellets were allowed to soften for 15 minutes and were vortexed for three minutes until the pellets were fully broken apart and homogeneous. Suspended bacteria were separated from fibrous material by centrifuging for five minutes (150 rpm). Supernatant was aliquoted into a sterile bottle and fecal slurries were pooled from multiple animals to create two separate solutions: male and female. The prepared bacterial solutions were placed in an anaerobic jar (BD BBL^TM^ GasPak^TM^ jar, Franklin Lakes, NJ, USA) and transported to the conventional vivarium. Each mouse received 150 μL donor material from the corresponding age- and sex-matched donor pooled sample. The remaining solution was frozen at -80°C. Mice received a single transplant and afterwards were placed in cages with dirty bedding from sex-matched Unaltered control donor mice.

#### DNA extraction of the fecal metagenome

Shotgun metagenomic sequencing was employed to determine the composition of the microbiota from fecal samples collected at 1, 3, 5, 13, 18, 18.5, 20, and 22 months of age. DNA extraction was conducted at the Benioff Center for Microbiome Medicine Microbial Genomics CoLab Plug-in at the University of California, San Francisco. Frozen fecal pellets were placed in 500 μL of cetyltrimethylammonium bromide (CTAB) extraction buffer (Lysing Matrix E tube, MP Biomedicals, Burlingame, CA, USA), vortexed and incubated (65°C, 15 min) (*50*). A mixture of 500 μL phenol, chloroform, and isoamyl alcohol (in a ratio of 25:24:1) was placed in the tube and the sample was subjected to bead-beating (5.5 m/s; 30 sec) and centrifuged (16,000 g, 5 min, 4°C). 400 μL of supernatant was placed in a 2 mL Eppendorf tube and 500 μL CTAB buffer was added. This process was repeated until the extraction volume reached 800 μL, after which 800 μL chloroform was added and centrifuged (3,000 g, 10 min). The supernatant (600 μL) was then placed in a 2 mL Eppendorf tube, 2-volume polyethylene glycol was added and the mixture was kept at 4°C overnight to allow for DNA precipitation. The samples were subsequently centrifuged (3,000 g, 60 mins), washed twice with chilled 70% ethanol, and placed in 100 μL of sterile water. Extracted DNA was quantified with Qubit 2.0 Fluorometer using dsDNA BR Assay Kit (Life Technologies, Grand Island, NY, USA).

#### Library preparation and quality control of fecal metagenome

Library preparation was performed at the Chan Zuckerberg Biohub San Francisco Genomics Platform. All materials and samples were processed in a post-PCR lab. Six 96-well plates containing extracted cDNA were quadrant pooled into two 384 deep-well plates, cDNA wells were normalized to 2 ng/μL and further diluted 1:10 to achieve a concentration of 0.2 ng/μL using SPT Labtech Firefly Liquid Handler. 0.5 μL of diluted cDNA was transferred to a regular 384 well plate using SPT Firefly and Integra ViaFLO384 liquid handlers, to be used as input for library preparation. Library preparation was completed using a modified Nextera XT DNA Library Kit protocol. This protocol has volumes that were miniaturized for high throughput processing in plate format. In summary, 0.5 μL (∼0.1 ng) of cDNA used as the input was tagmented at 55°C for 5 minutes using Tagment DNA Buffer (0.2 μL) and Amplicon Tagment mix (1.2 μL). The tagmentation reaction was neutralized by adding 0.4 μL of Neutralize Tagment Buffer (NT) followed by 2000 RCF centrifugation for 5 minutes. Index PCR was performed by adding 1.2 μL of Nextera PCR mix (NPM), 0.4 μL of 5 μM i5 and i7 indexing primers while ensuring each library received a unique combination of i5 and i7. Libraries were amplified according to the following cycling parameters: 72°C for 3 min; 95°C for 30 sec; 12 cycles (95°C for 10 sec, 55°C for 30 sec, 72°C for 30 sec) and final extension at 72°C for 5 min. Barcoded libraries were pooled to a 1.5 mL lo-bind Eppendorf tube using SPT Labtech MosquitoLV. Pooled libraries were processed through two 0.7 X SPRI cleanups and were quality controlled for sequencing using Aligent 4150 Tapestation System and D5000 ScreenTape and via qPCR using Bio-Rad CFX96 RT System. All libraries were pooled equimolar for loading on a Novaseq 6000 instrument.

#### Shotgun metagenomic sequencing of fecal metagenome

Sequencing was performed at the Chan Zuckerberg Biohub San Francisco Genomics Platform. All libraries were sequenced using the loading guidelines provided by Illumina. The Illumina sequencing library was loaded with a NovaSeq 6000 instrument using two S4 kits (2 x 10 billion reads) flow cells using PE150 run parameter.

#### Quality control, taxonomic profiling metagenomes, and antimicrobial resistance gene identification

The open-source CZ ID pipeline (https://czid.org/) was used for quality control processing, the taxonomic classification of microbes (mNGS pipeline version 8.3), and antimicrobial resistance genes (AMR pipeline version 1.4.2; CARD Database version 3.2.6; Wildcard Database version 4.0.0). Microbial identification in the CZ ID pipeline involved subtractive alignment to mouse host genome (Bowtie2) using raw FASTQ files and fastq quality control (*2*). Microbial reads were aligned to the NCBI NT database (National Center for Biotechnology Information Nucleotide Database) using minimap2 (assembly-based alignment) (*2*). Microbial reads were also aligned to the NCBI NR database (non-redundant protein) (*2*). Prior to quality control processing samples had an average number of 62,212,731± 30,371,926 reads. Samples that had more than 150,000 reads were kept for downstream microbial and antimicrobial resistance genes analyses. After quality control, samples had an average number of 30,557,604 ± 13,488,671 reads (table S9). Following taxonomic alignment, samples were run through CZ ID’s antimicrobial resistance pipeline (Comprehensive Antibiotic Resistance Database Resistance Gene Identifier tool) and antimicrobial resistance genes that had ≥ 5% read coverage breadth were kept for resistance analyses (*2*). A table of identified antimicrobial resistance genes and their microbial origin and resistance classification are provided (table S10).

#### Flow cytometry immune profiling

Spleens were collected at euthanasia in individual tubes on ice with 5 mL complete RMPI medium, which contained 10% fetal bovine serum, 2 mM L-glutamine, 1% penicillin-streptomycin (Corning), 2.5 μg/mL amphotericin B (Gibco), 1 mM sodium pyruvate (Corning), and 10 mM HEPES (Corning). The spleens were processed through a 70 μm cell strainer with isolation buffer comprising PBS supplemented with 2% fetal bovine serum, 1% penicillin-streptomycin, and 1 mM EDTA. The resulting single-cell suspension was centrifuged at 350×g for five minutes at room temperature. The pellet was then resuspended in 1 mL of ACK lysis buffer (Thermo Fisher Scientific) and incubated for three minutes. Additional isolation buffer was added to adjust the total volume to 10 mL, followed by another centrifugation. After washing in isolation buffer, the cells were resuspended in 1 mL of freezing medium consisting of 90% fetal bovine serum and 10% DMSO, and stored in a liquid nitrogen tank until subsequent thawing for analytical procedures.

Frozen splenocytes were thawed and analyzed for T cell, B cell, and myeloid populations using multicolor spectral flow cytometry. All antibodies were used at a 1:200 dilution in FACS buffer (PBS with 2% FBS and 1 mM EDTA). Dead cells were excluded using Zombie UV viability dye. The specific staining panels are detailed below:

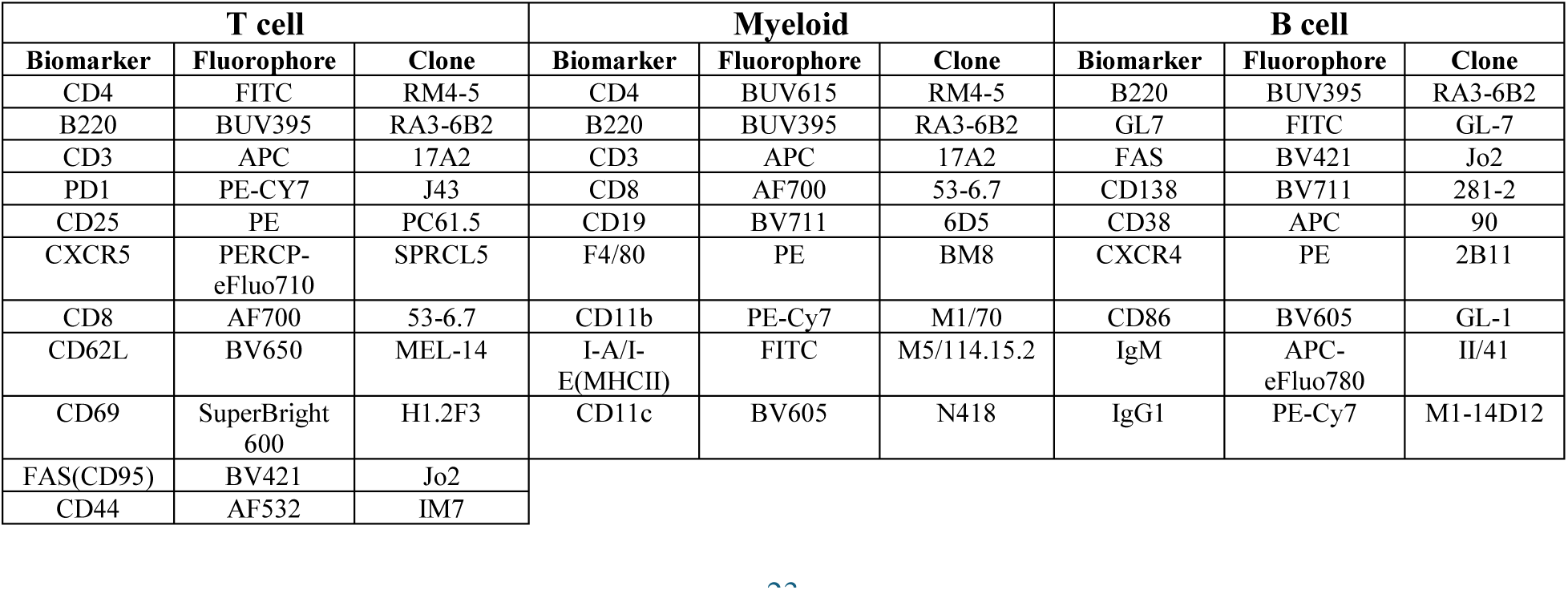
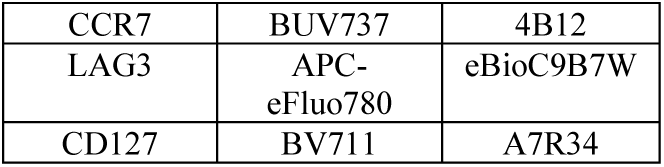

| T cell |  |  | Myeloid |  |  | B cell |  |  |
| --- | --- | --- | --- | --- | --- | --- | --- | --- |
| Biomarker | Fluorophore | Clone | Biomarker | Fluorophore | Clone | Biomarker | Fluorophore | Clone |
| CD4 | FITC | RM4-5 | CD4 | BUV615 | RM4-5 | B220 | BUV395 | RA3-6B2 |
| B220 | BUV395 | RA3-6B2 | B220 | BUV395 | RA3-6B2 | GL7 | FITC | GL-7 |
| CD3 | APC | 17A2 | CD3 | APC | 17A2 | FAS | BV421 | Jo2 |
| PD1 | PE-CY7 | J43 | CD8 | AF700 | 53-6.7 | CD138 | BV711 | 281-2 |
| CD25 | PE | PC61.5 | CD19 | BV711 | 6D5 | CD38 | APC | 90 |
| CXCR5 | PERCP-eFluo710 | SPRCL5 | F4/80 | PE | BM8 | CXCR4 | PE | 2B11 |
| CD8 | AF700 | 53-6.7 | CD11b | PE-Cy7 | M1/70 | CD86 | BV605 | GL-1 |
| CD62L | BV650 | MEL-14 | I-A/I-E(MHCII) | FITC | M5/114.15.2 | IgM | APC-eFluo780 | II/41 |
| CD69 | SuperBright 600 | H1.2F3 | CD11c | BV605 | N418 | IgG1 | PE-Cy7 | M1-14D12 |
| FAS(CD95) | BV421 | Jo2 |  |  |  |  |  |  |
| CD44 | AF532 | IM7 |  |  |  |  |  |  |

|  |  |  |
| --- | --- | --- |
| CCR7 | BUV737 | 4B12 |
| LAG3 | APC-eFluo780 | eBioC9B7W |
| CD127 | BV711 | A7R34 |

#### Gut permeability histology

Fresh colon tissue was harvested from mice and stored at -80°C before processing for RNA extraction using Trizol and cDNA synthesis. Real-time quantitative PCR was performed using the QuantStudio 7 Real-Time PCR system using TaqMan Universal PCR Master Mix and TaqMan Gene Expression Assays for the following genes: *ppia* (housekeeping), *cldn2*, *occluding*, *tip1*, and *muc2* (Applied Biosystems). Relative expression of transcript levels were measured using the 2^-ΔΔCt^ method using amplified Ct values.

#### Serum/plasma biomarker content

Whole blood was collected using cardiac puncture at euthanasia under anesthesia. For serum collection, whole blood was placed in sterile 1.5 mL microcentrifuge tubes, allowed to clot, centrifuged for 10 minutes, and placed in 25 μL aliquots and frozen at -80°C. For plasma collection, whole blood was placed in EDTA (ethylenediaminetetraacetic acid)-coated microcentrifuge tubes (BD Microtainer® Blood Collection Tubes, 365992, Franklin Lakes, NJ, USA), centrifuged at 4°C for 10 minutes, and placed in 25 μL aliquots and frozen at -80°C. An ELISA (enzyme-linked immunosorbent assay) was carried out at Duke Molecular Physiology Institute Biomarkers Shared Resource. Serum was collected from fasted mice and was measured for adiponectin, leptin, and insulin. Plasma was collected from unfasted mice and was measured for estrogen (females only), interleukin 6 (IL-6), tumor necrosis factor-α (TNF-α), and insulin-like growth factor 1 (IGF-1).

#### Statistics

Principal coordinate analyses (PCoA) were performed on rarefied Bray-Curtis dissimilarity matrices on ARGs and taxa to discern differences in composition amongst treatment groups, timepoints, and cage assignments. Differences in Shannon diversity, body weight, fat pad mass, and serum/plasma biomarkers were evaluated using a one-way analysis of variance (ANOVA) with a post hoc Dunnett test to evaluate significance between each treatment group relative to the Unaltered sex-matched control. Correlations between individual ARGs and microbial species were determined using a pairwise Spearman correlation calculation followed by a Benjamini-Hochberg FDR correction. To determine ARGs and genera that significantly changes with time within individual treatment groups by sex Microbiome Multivariate Associations with Linear Models (MaAsLin2) was performed in which cage assignment was accounted for as a random effect. Post hoc Benjamini-Hochberg FDR q-values were reported along with p-values. Taxa heatmaps were generated by averaging the relative abundance of 15 top taxa within each timepoint from the Continuous Dose group. A pseudocount of 0.1 was used and the fold change of the relative abundance of each taxa was calculated relative to the baseline value at 1 month of age with a log2 transform. Here, we defined colonization as the detection of genera that were not observed prior to antibiotic exposure (less than 0.1% relative abundance) (*12*). Colonizers were not detected prior to antibiotic exposure and remained at an increased abundance with prolonged antibiotic exposure. Dynamic colonizers were not detected prior to antibiotic exposure and exhibited temporary increases or decreases in abundance with prolonged antibiotic exposure. Genera that changed in abundance were present prior to antibiotic exposure and remained increased or decreased with prolonged antibiotic exposure. Genera with a dynamic change in abundance were present prior to antibiotic exposure and exhibited temporary increases or decreases in abundance with prolonged antibiotic exposure. ARG heatmaps were generated by averaging the relative abundance of top ARGs within each timepoint from the Continuous Dose group with a log2 transform. Variance of % ARG abundance associated with *Enterobacteriaceae*, ARG richness, ARG abundance, and relative abundance of taxa was calculated and followed with Shapiro’s test to confirm normality and a Likelihood Ratio Test was used to compare two Generalized Least Squares models with homoscedastic and heteroscedastic residual variance to determine changes in variance over time. Rate constants associated with changes in taxonomy in the first four months after the start of antibiotic dosing were determined with regression models. All statistical testing was completed using R Statistical Software (v 4.0.3; R Core Team 2020).

**Fig. S1.**
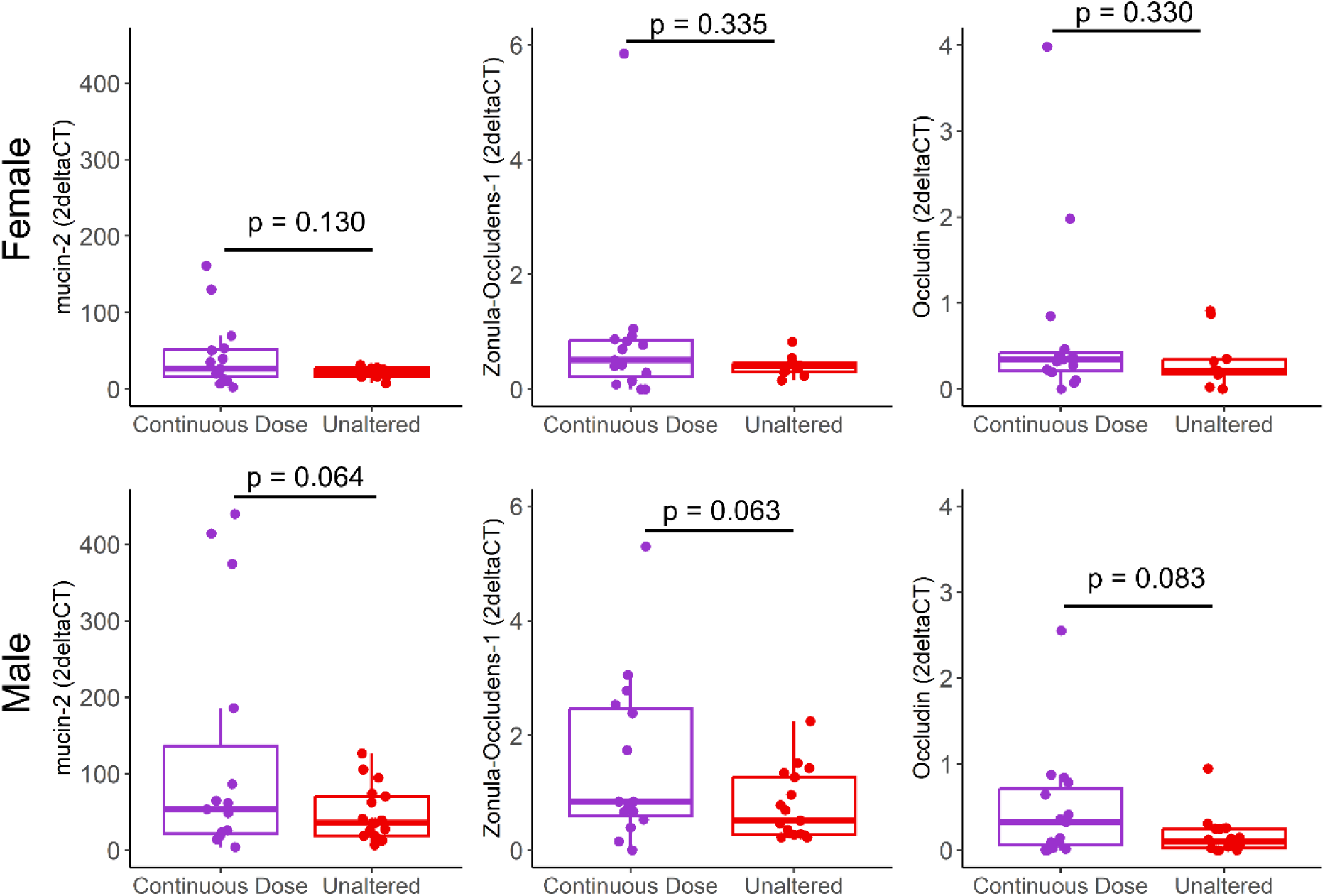
Gut permeability for Unaltered and Continuous Dose mice. Continuous antibiotic exposure did not significantly influence gut permeability.

**Fig. S2.**
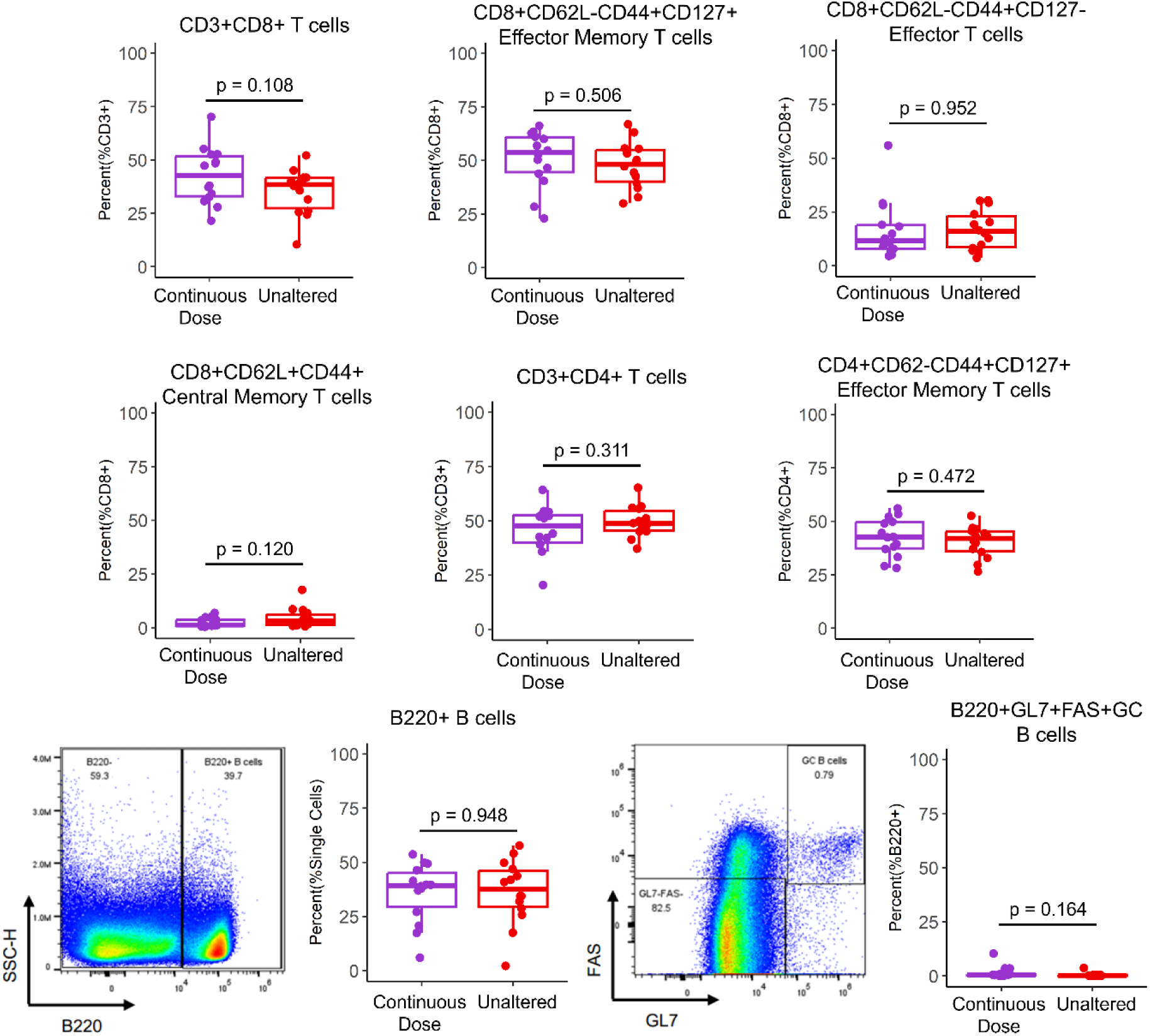
Splenic immune cell profiling for Unaltered and Continuous Dose mice. Continuous antibiotic exposure did not significantly influence splenic immune cell profile.

**Fig. S3.**
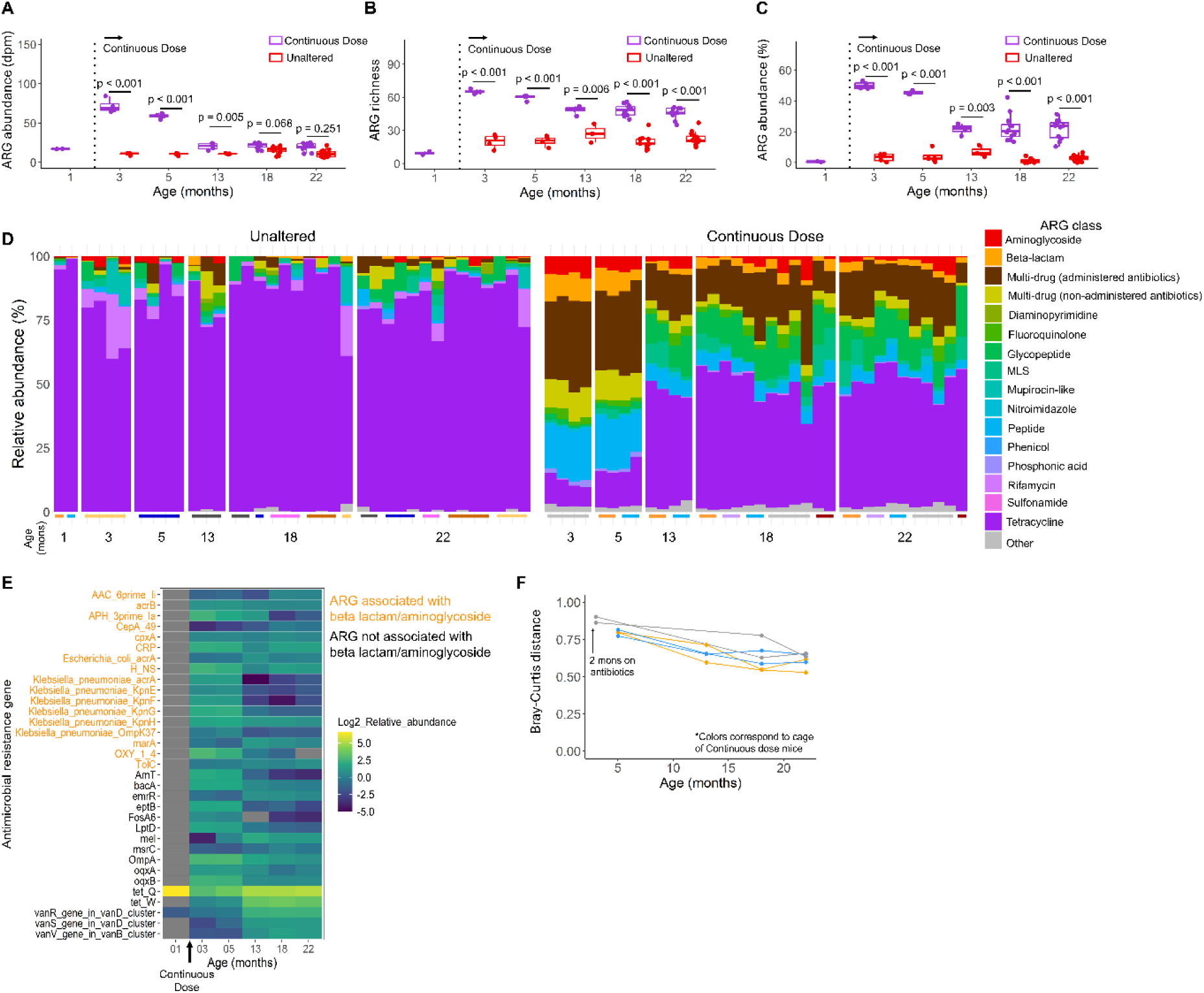
Effects of Continuous exposure to antibiotics on the resistome of male mice. (A) In the first few months of antibiotic dosing there was an increase in ARG abundance that peaked at three months of age, however, the abundance of ARGs declined and reached levels indistinguishable from that of the Unaltered group by 18 months of age. (B) In the first few months of antibiotic dosing there was an increase in ARG richness that peaked at three months of age and declined shortly after but remained increased relative to the Unaltered group for the remainder of life. (C) Two months after the start of antibiotic exposure < 50% of the ARG burden was attributed to resistance to the classes of antibiotics that were administered (beta-lactams and aminoglycosides). (D) Continuous exposure to antibiotics resulted in dynamic changes in the resistome composition. (E) The increase in ARG richness was primarily due to the sustained presence of ARGs that were first detected shortly after the start of antibiotic dosing. (F) Longitudinal tracking in individual mice showed that Continuous antibiotic exposure resulted in changes in the Bray-Curtis resistome composition over time that shifted towards the baseline Unaltered composition. Lines connect data points from the same individual and represent Bray-Curtis distance to baseline (one month of age prior to antibiotic exposure).

**Fig. S4.**
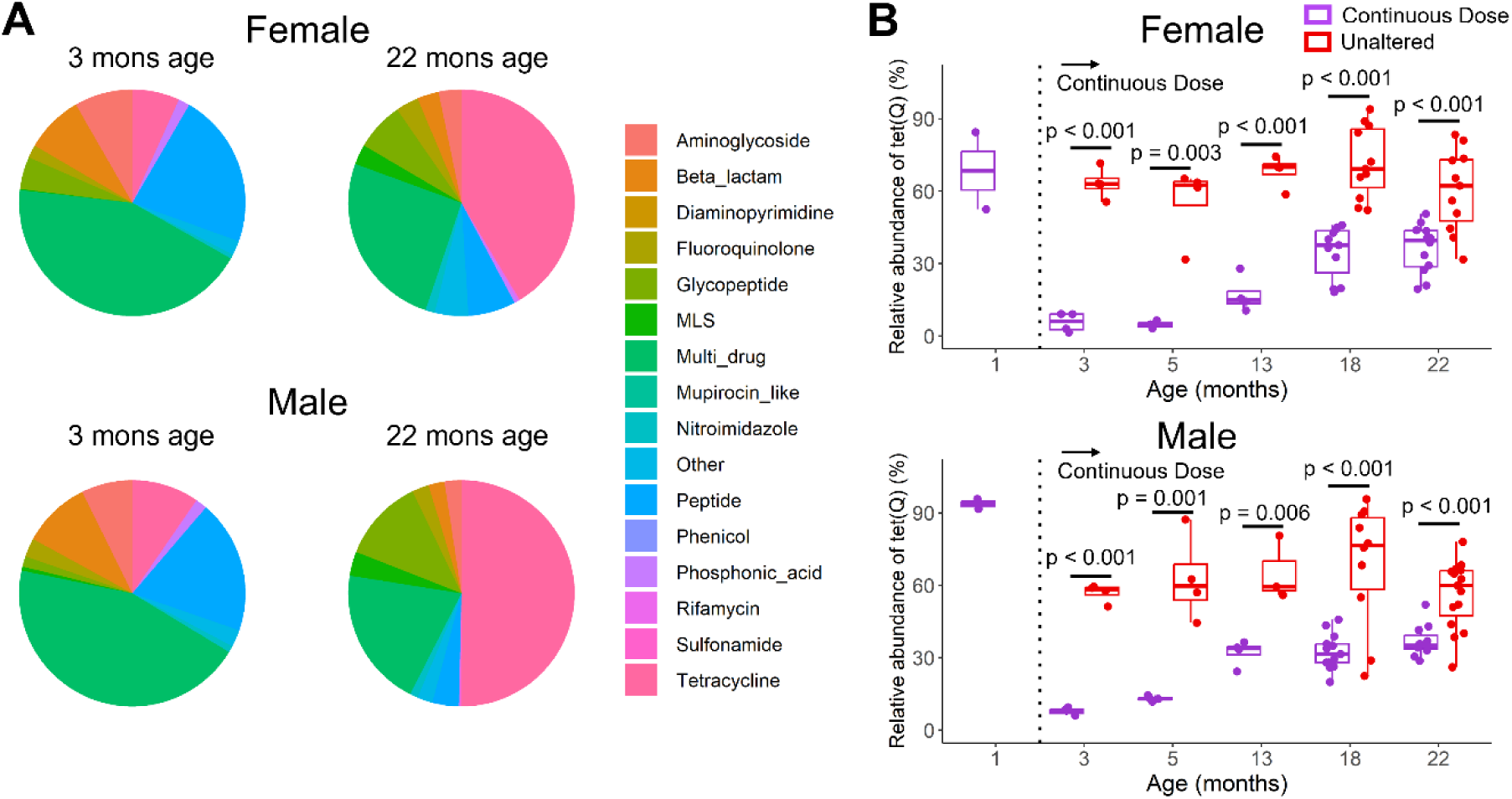
(A) Average abundance of ARGs associated with different classes of antibiotics in Continuous Dose mice revealed a decrease in ARGs associated with multidrug resistance and increase in ARGs associated with tetracycline over time. (B) The relative abundance of *tet(Q)* constituted the majority of the resistome in mice prior to antibiotic exposure, was significantly decreased with antibiotic exposure, and gradually increased with prolonged antibiotic exposure.

**Fig. S5.**
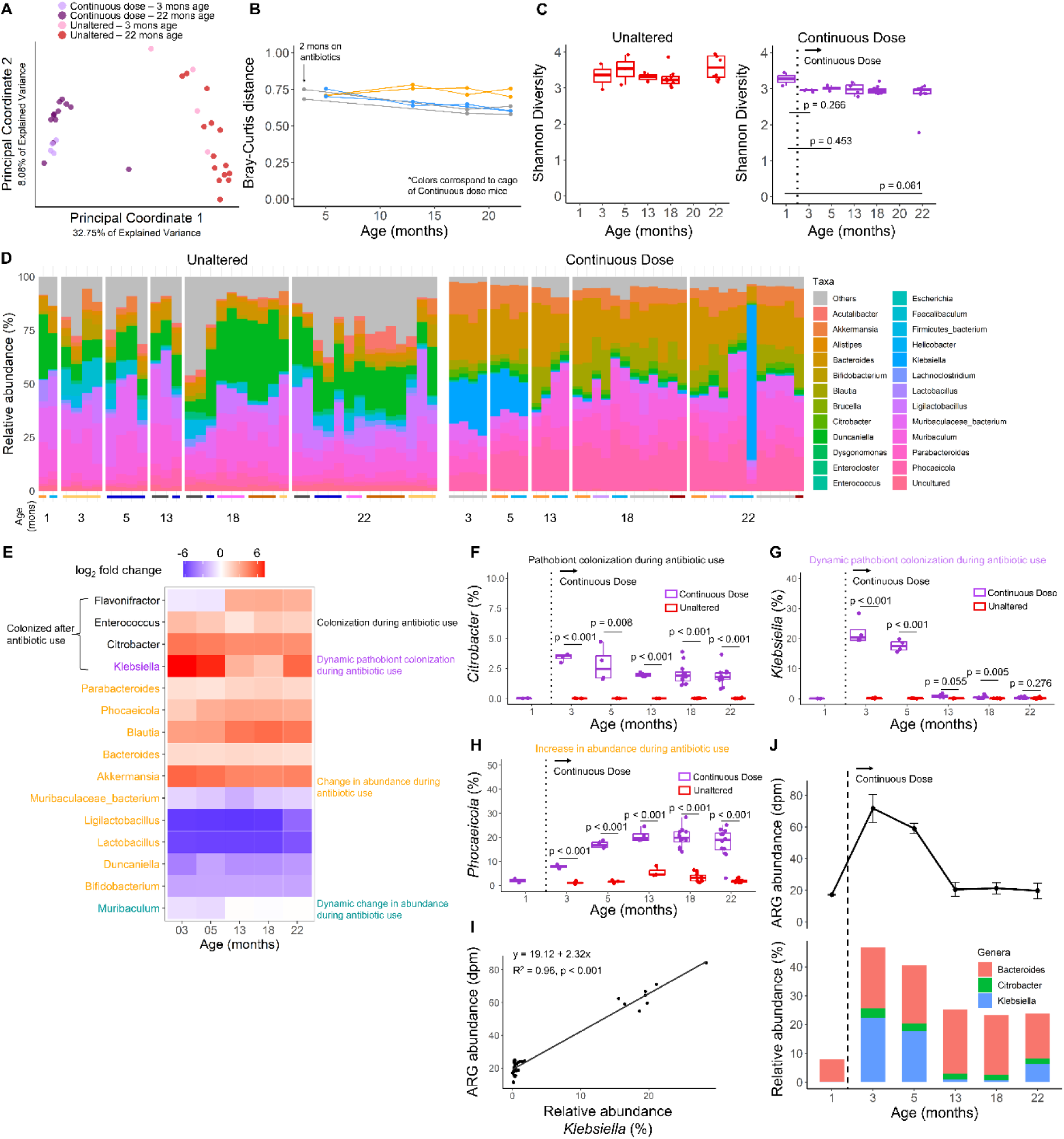
Effects of Continuous exposure to antibiotics on the microbiota of male mice. (A-B) Continuous exposure to oral antibiotics resulted in a rapid, drastic change in gut microbial composition compared to mice without antibiotic exposure and composition changed over time. (C) Shortly after the start of Continuous antibiotic dosing the Shannon diversity of the taxa decreased. (D-E) Genera-level analysis revealed complex dynamics of taxa during antibiotic dosing. (F-H) Representative plots in males with relative abundance of genera classified based on their response to antibiotics. (I) Regression of the relative abundance of *Klebsiella* against ARG abundance in Unaltered and Continuous Dose mice demonstrated that changes in ARG abundance were strongly linked with abundance of *Klebsiella* (Pearson correlation: 0.98). (J) Shifts in relative abundance of ARG harboring taxa directly align with shifts in ARG abundance.

**Fig. S6.**
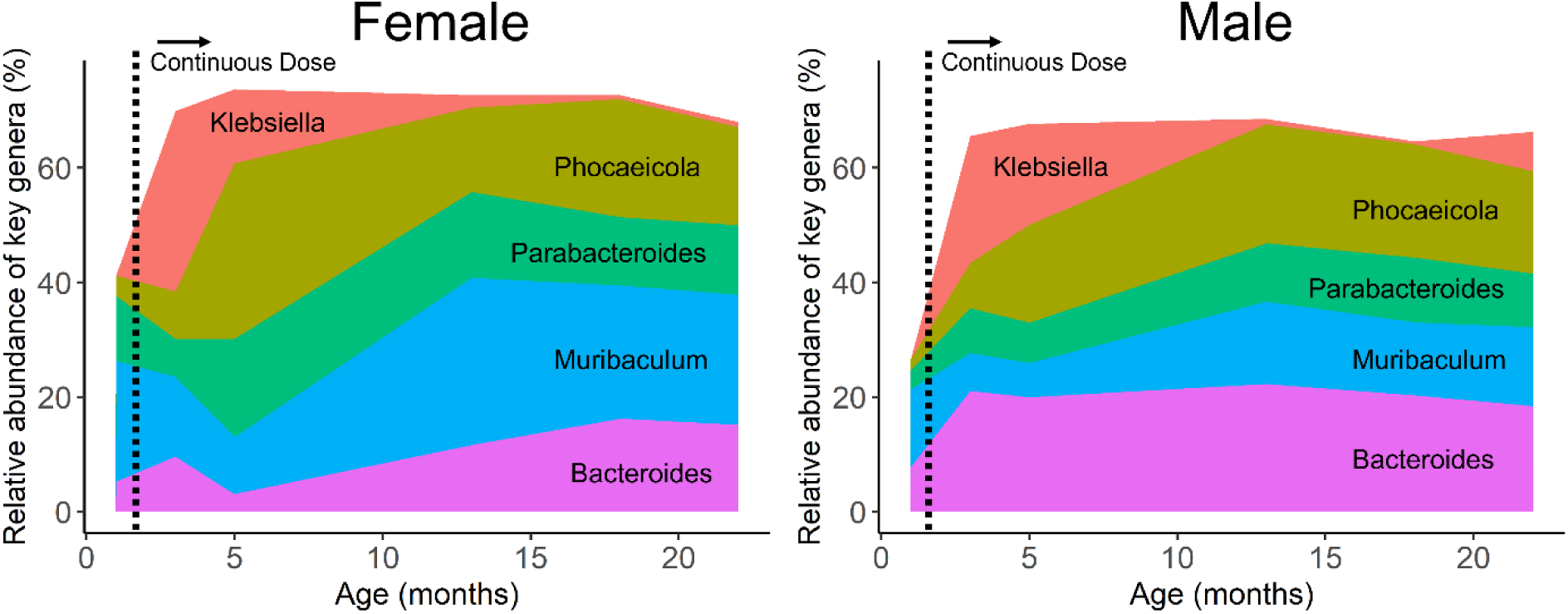
Average relative abundance of key genera (*Bacteroides*, *Klebsiella*, *Muribaculum*, *Parabacteroides*, *Phocaeicola*) over time in Continuous Dose mice revealed a dynamic shift amongst microbes.

**Fig. S7.**
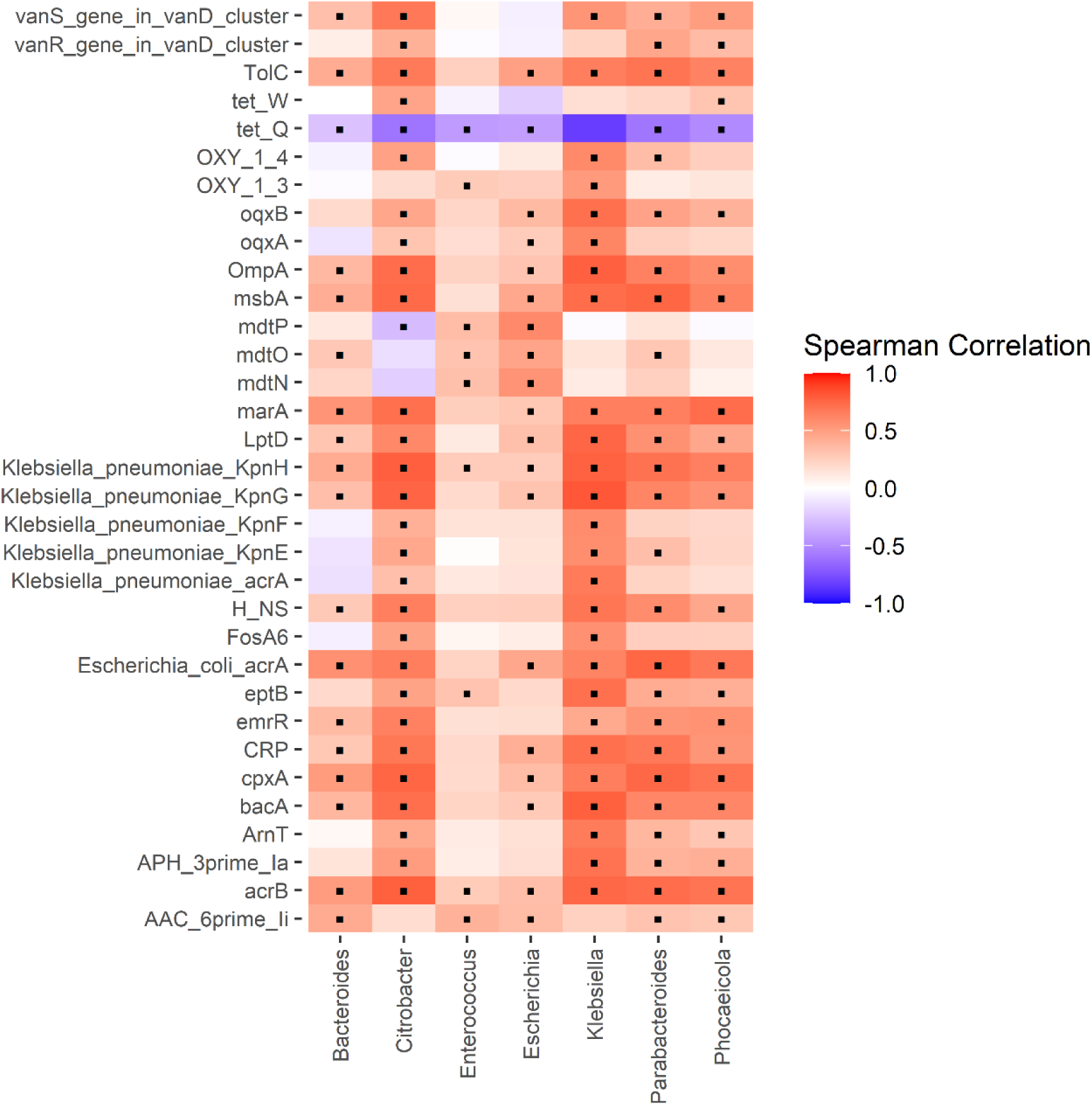
Pairwise Spearman correlation between genera and ARGs in Continuous Dose female mice; ▪ indicates a significant correlation (p < 0.05, Benjamini-Hochberg correction FDR).

**Fig. S8.**
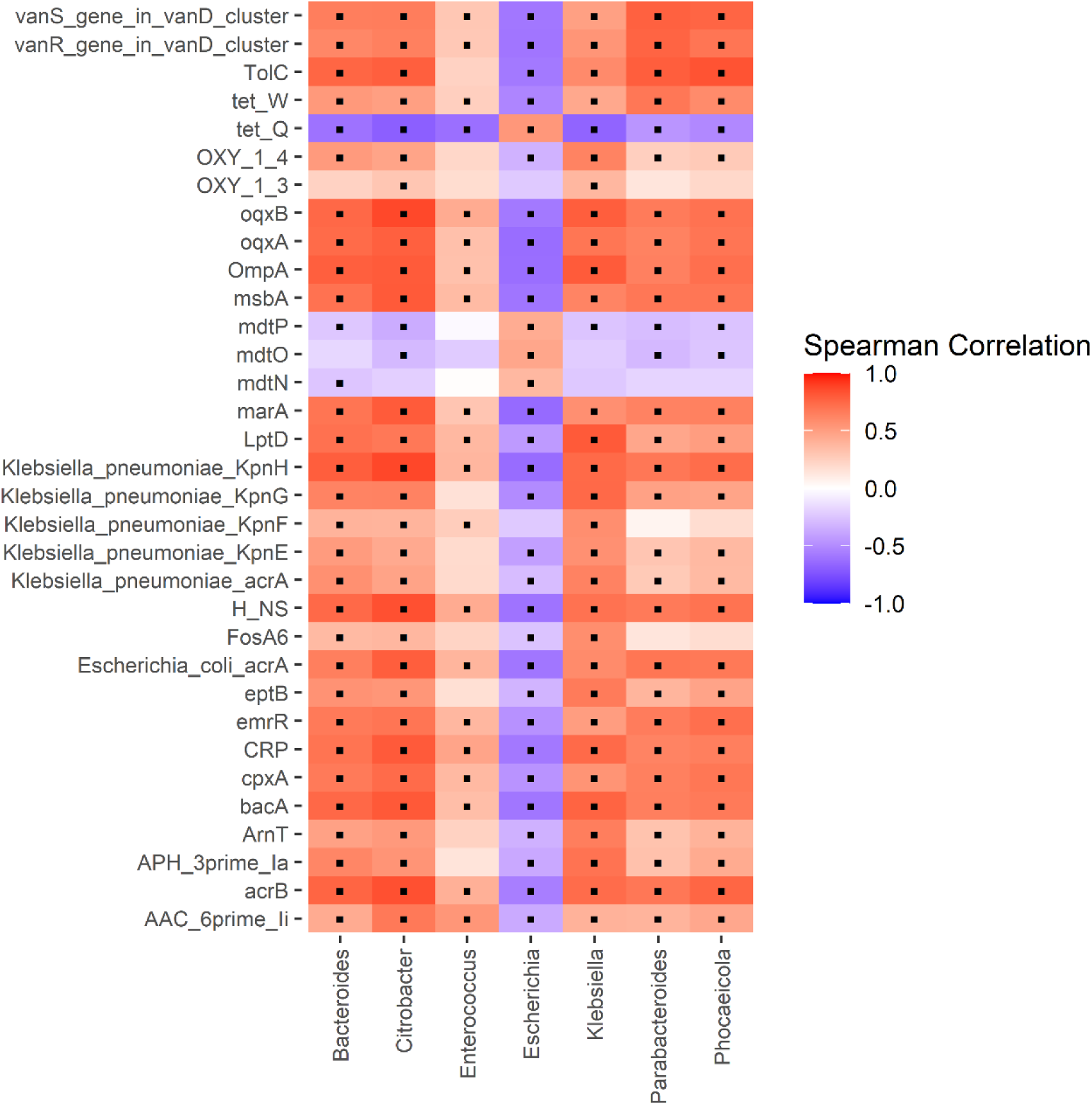
Pairwise Spearman correlation between genera and ARGs in Continuous Dose male mice; ▪ indicates a significant correlation (p < 0.05, Benjamini-Hochberg correction FDR).

**Fig. S9.**
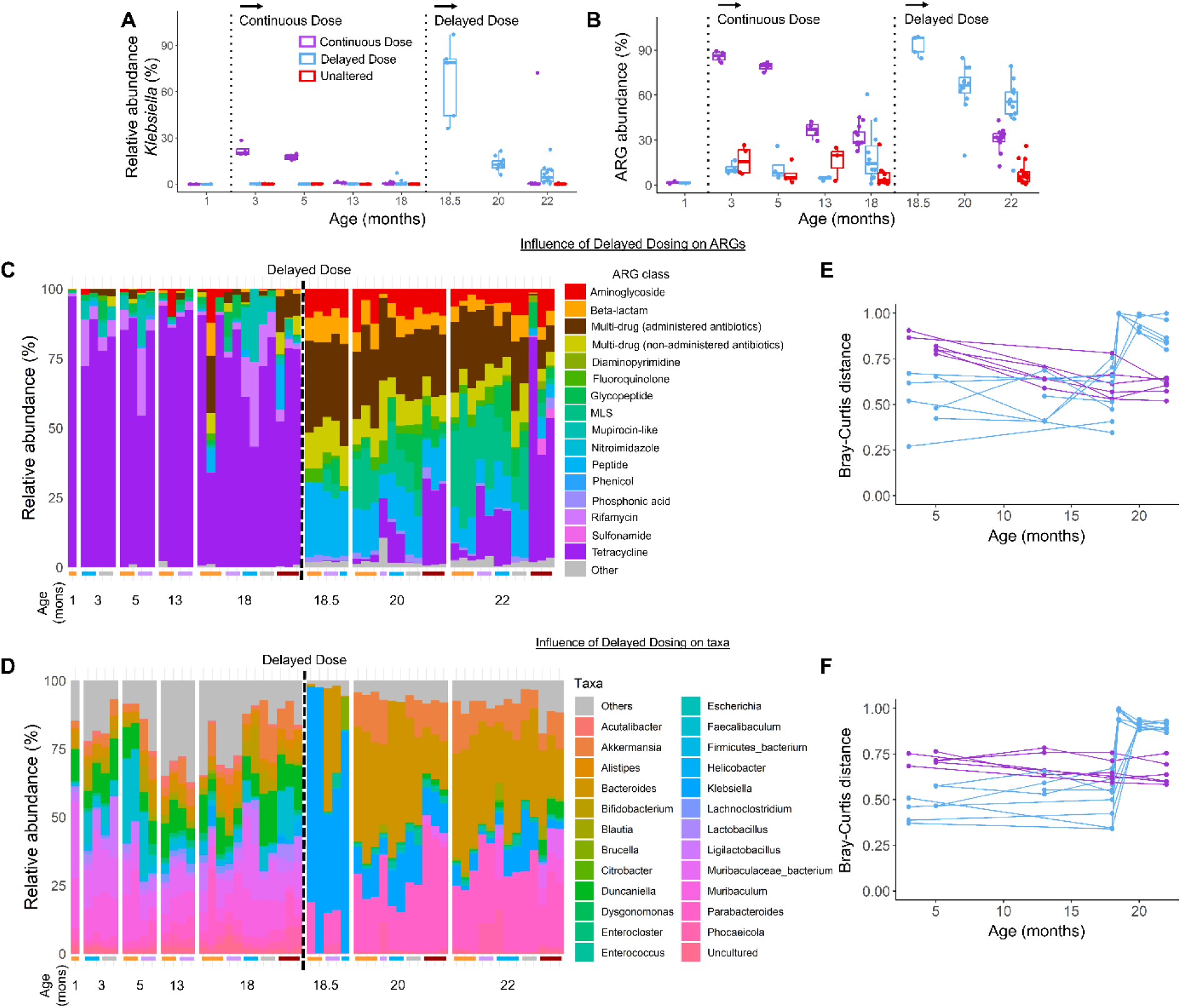
Effects of Delayed Dose antibiotic exposure on the microbiota and resistome of male mice. (A) Mice in the Delayed Dose group had a transient increase in *Klebsiella* and more rapid decline in *Klebsiella* relative to mice in the Continuous Dose group that were exposed to antibiotics for the same duration. (B) Delayed Dose mice had a transient increase in the relative abundance of ARGs that were associated with *Enterobacteriaceae*. (C-D) Composition of the resistome and microbiota over time in individual mice. (E-F) Bray-Curtis distance from baseline in the resistome and microbiota composition of Delayed Dose male mice.

**Fig. S10.**
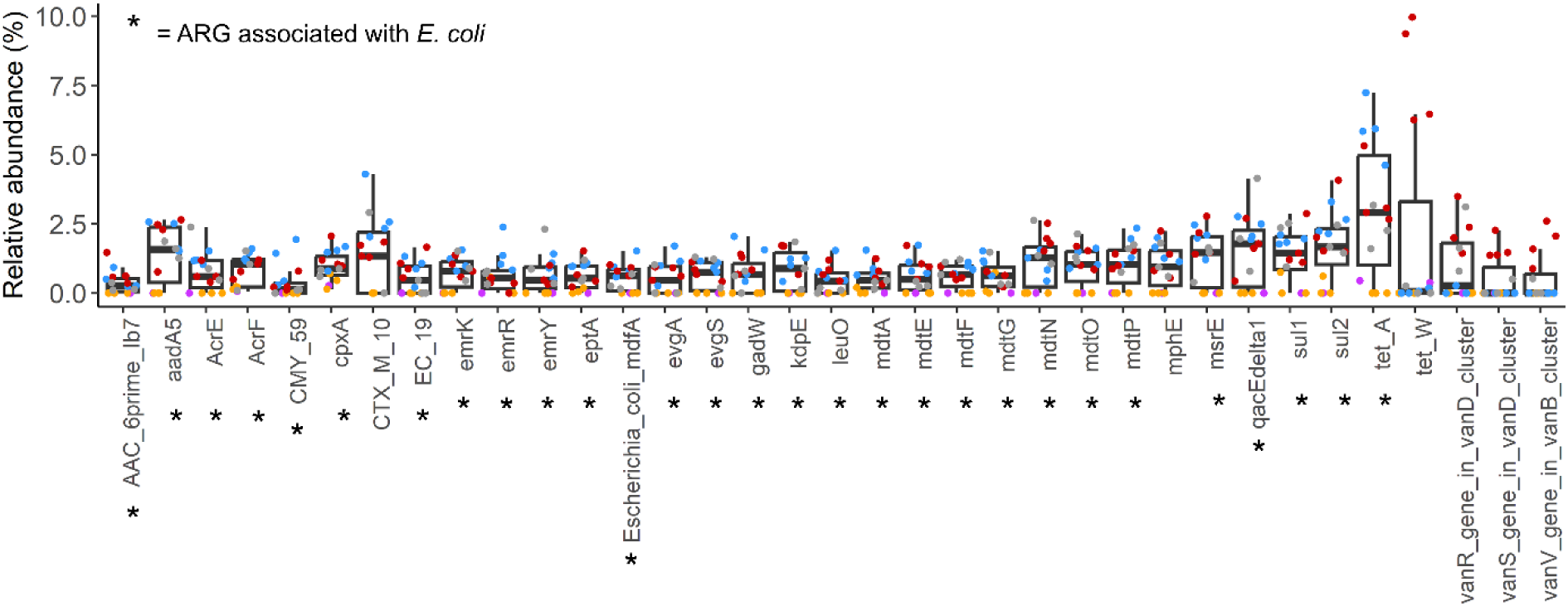
Outliers in the relationship between *Klebsiella* abundance and ARG burden in Figure 3F had increased relative abundance of more than 29 ARGs associated with *E. coli* and more than 5 ARGs associated with other taxa. Colored dots indicate mice housed in the same cage.

**Fig. S11.**
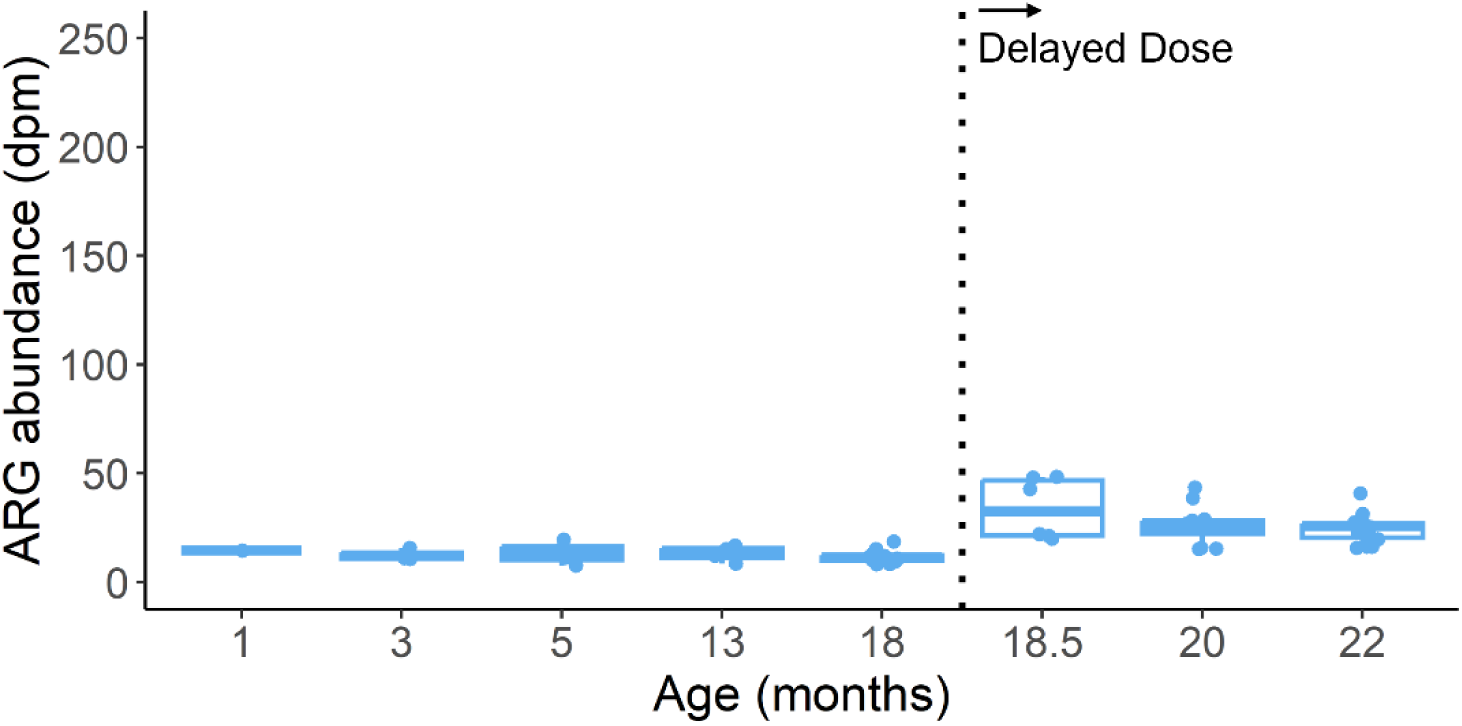
After subtracting off contributions of ARGs associated with *E. coli* from total ARG abundance in Delayed Dose female mice, ARG abundance declined after exposure to antibiotics as observed in male mice.

**Fig. S12.**
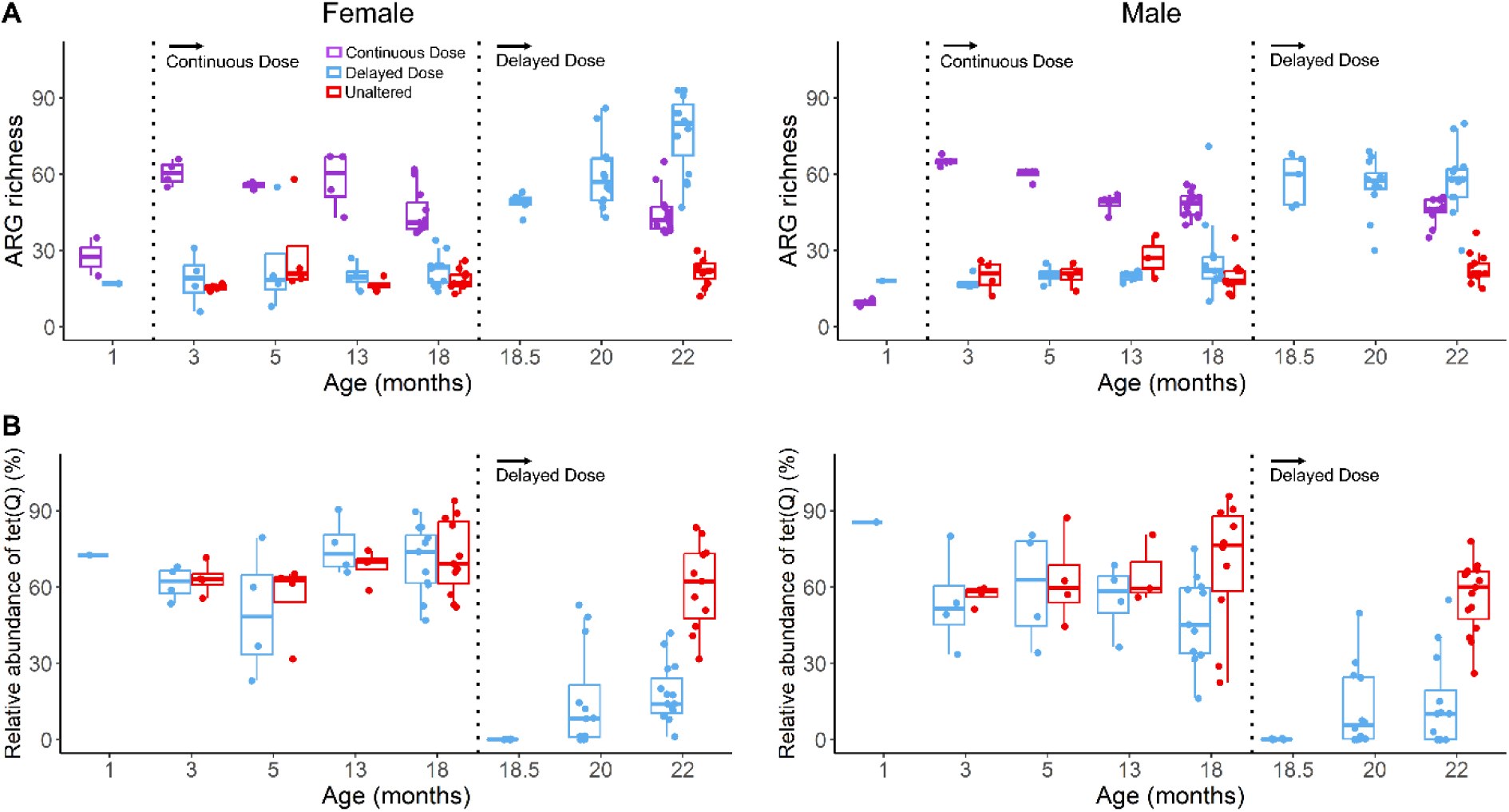
(A) ARG richness in Delayed Dose, Continuous Dose, and Unaltered groups over time. (B) *tet(Q)* dominated the resistome prior to antibiotic exposure and slowly increased in abundance after prolonged antibiotic exposure.

**Fig. S13.**
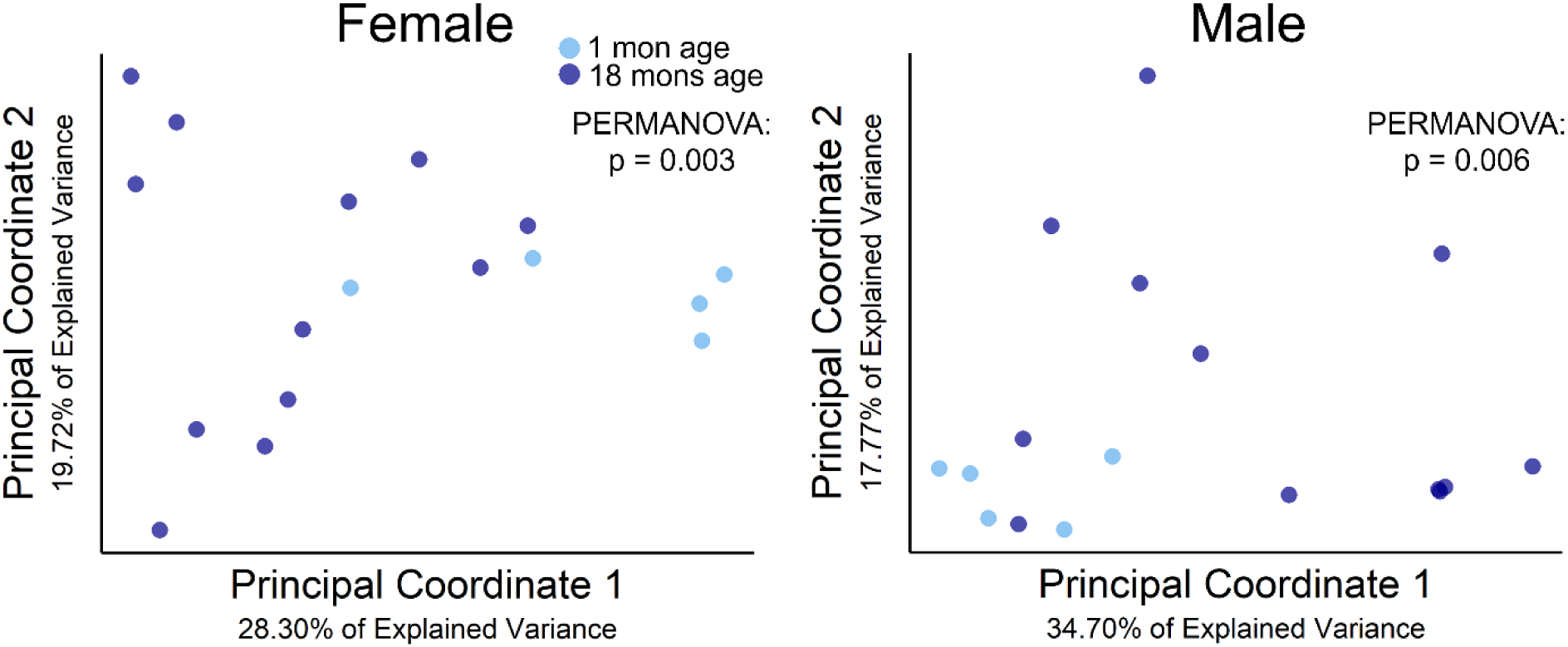
Taxonomic composition of the gut microbiota in Delayed Dose mice significantly shifted over aging from one to 18 months of age in both females and males.

**Fig. S14.**
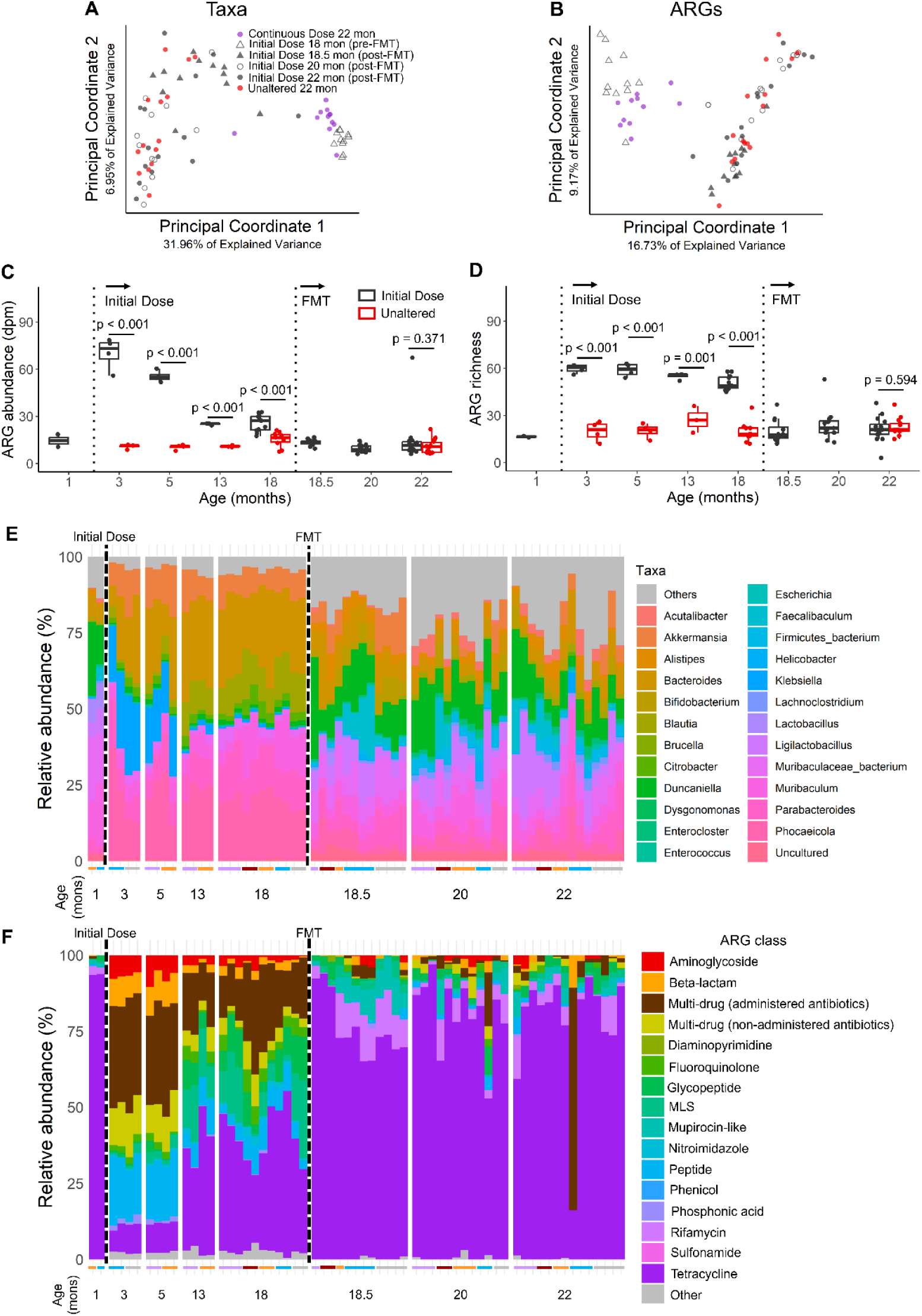
(A-B) As observed in female mice, male mice that received the FMT after prolonged antibiotic exposure had their microbiota composition and resistome restored to the Unaltered state. (C-D) As observed in female mice, male mice that receive the FMT after prolonged antibiotic exposure had their total ARG burden and richness restored to the Unaltered state. (E-F) Composition of the microbiota and resistome prior to and after receiving the FMT.

**Fig. S15.**
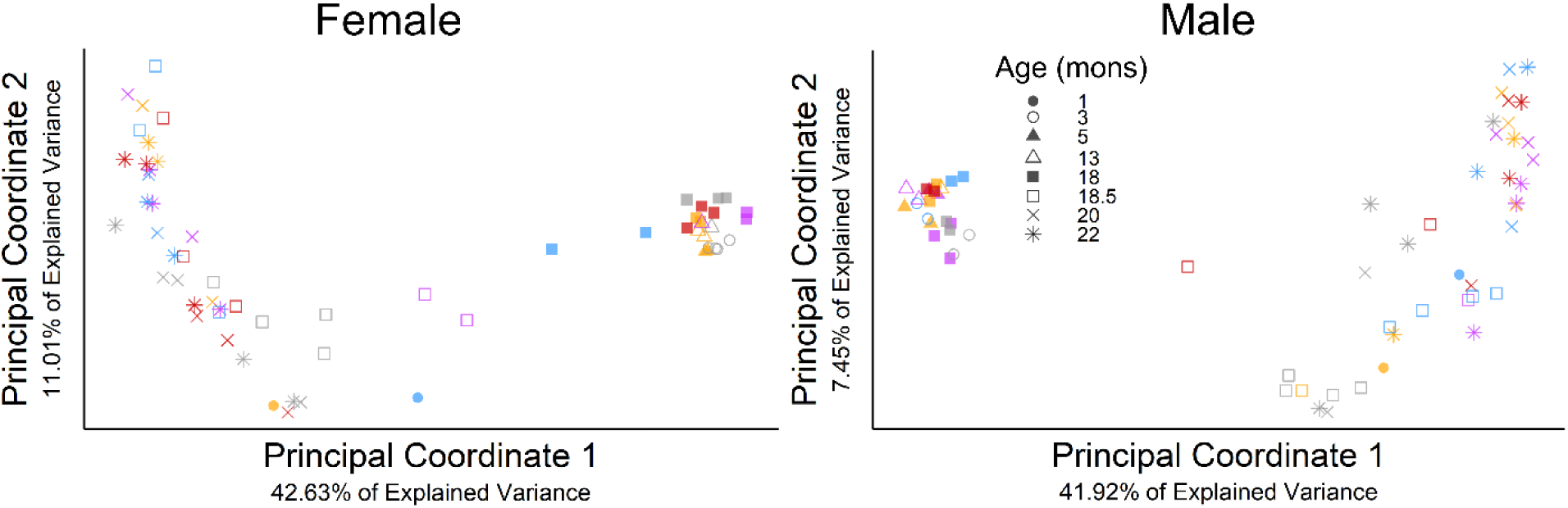
Longitudinally tracking the Bray-Curtis distance principal coordinate analysis of the Initial Dose group.

**Table S1.** Circulating fasted serum (adiponectin, leptin, insulin) and non-fasted plasma (IL-6, IGF-1, TNF-α, estrogen – female only) biomarkers. Average values with standard deviation are shown. Dunnett test performed relative to Unaltered sex-matched control.

| Treatment Group | Adiponectin (ng/mL) | Leptin (pg/mL) | Insulin (pg/mL) | IL-6 (pg/mL) | IGF-1 (pg/mL) | TNF-α (pg/mL) | Estrogen (pg/mL) |
| --- | --- | --- | --- | --- | --- | --- | --- |
| *Female* |  |  |  |  |  |  |  |
| Unaltered | 11791.600 ± 1854.456 | 6664.927 ± 4513.005 | 328.640 ± 284.080 | 59.407 ± 40.367 | 323744.286 ± 30842.775 | 25.336 ± 17.843 | 739.319 ± 414.776 |
| Continuous Dose | 11603.600 ± 1316.444<br>p = 0.994 | 3155.279 ± 2115.732<br>p = 0.012 | 233.815 ± 229.871<br>p = 0.516 | 95.657 ± 94.067<br>p = 0.612 | 264990.000 ± 34082.415<br>p = 0.007 | 20.651 ± 5.155<br>p = 0.998 | 1240.004 ± 892.425<br>p = 0.204 |
| Delayed Dose | 11601.000 ± 1365.696<br>p = 0.993 | 3532.141 ± 1990.284<br>p = 0.027 | 197.989 ± 113.136<br>p = 0.268 | 70.281 ± 67.956<br>p = 0.980 | 293688.571 ± 42610.087<br>p = 0.226 | 25.148 ± 15.417<br>p = 1.000 | 833.631 ± 274.280<br>p = 0.974 |
| Initial Dose | 10307.100 ± 1173.284<br>p = 0.281 | 3160.301 ± 1975.363<br>p = 0.012 | 202.444 ± 107.547<br>p = 0.294 | 102.700 ± 48.065<br>p = 0.481 | 359667.143 ± 17646.433<br>p = 0.123 | 74.361 ± 114.718<br>p = 0.295 | 640.484 ± 239.664<br>p = 0.971 |
| *Male* |  |  |  |  |  |  |  |
| Unaltered | 6399.000 ± 304.501 | 3374.644 ± 2181.318 | 290.774 ± 186.042 | 87.982 ± 64.059 | 293718.571 ± 23523.576 | 11.359 ± 3.214 | -- |
| Continuous Dose | 6002.960 ± 935.092<br>p = 0.834 | 2283.911 ± 1341.219<br>p = 0.219 | 327.804 ± 197.484<br>p = 0.883 | 48.077 ± 29.648<br>p = 0.236 | 288605.714 ± 20158.671<br>p = 0.979 | 16.119 ± 5.251<br>p = 0.180 | -- |
| Delayed Dose | 5962.880 ± 1396.125<br>p = 0.794 | 2099.470 ± 1129.790<br>p = 0.126 | 238.974 ± 99.210<br>p = 0.743 | 73.761 ± 39.391<br>p = 0.876 | 295170.000 ± 27974.134<br>p = 0.999 | 8.669 ± 5.436<br>p = 0.599 | -- |
| Initial Dose | 5274.200 ± 654.134<br>p = 0.165 | 2277.777 ± 1359.676<br>p = 0.215 | 242.443 ± 99.685<br>p = 0.779 | 63.823 ± 34.160<br>p = 0.611 | 298610.000 ± 44024.678<br>p = 0.981 | 13.854 ± 4.967<br>p = 0.650 | -- |

**Table S2.** Body weight at euthanasia and perigonadal fat pad mass. Average values with standard deviations are shown. Dunnett test performed compared to Unaltered sex-matched control.

| Treatment group | Body weight at euthanasia (g) | Perigonadal fat pad mass (g) |
| --- | --- | --- |
| *Female* |  |  |
| Unaltered | 27.343 ± 4.863 | 0.773 ± 0.386 |
| Continuous Dose | 24.194 ± 2.073<br>p = 0.015 | 0.452 ± 0.180<br>p = 0.003 |
| Delayed Dose | 24.356 ± 2.619<br>p = 0.025 | 0.495 ± 0.235<br>p = 0.014 |
| Initial Dose | 24.353 ± 2.149<br>p = 0.022 | 0.443 ± 0.203<br>p = 0.002 |
| *Male* |  |  |
| Unaltered | 30.094 ± 2.180 | 0.621 ± 0.207 |
| Continuous Dose | 28.500 ± 2.384<br>p = 0.176 | 0.461 ± 0.182<br>p = 0.154 |
| Delayed Dose | 31.239 ± 3.735<br>p = 0.432 | 0.635 ± 0.422<br>p = 0.997 |
| Initial Dose | 29.710 ± 2.115<br>p = 0.944 | 0.512 ± 0.156<br>p = 0.424 |

**Table S3.** ARGs that significantly changed with time (3-22 months) in the Continuous Dose group with an abundance greater than 1%.

| ARG | Change over time | p-value | q-value |
| --- | --- | --- | --- |
| *Female* |  |  |  |
| *Klebsiella pneumoniae*<br>*acrA* | Decrease | < 0.001 | < 0.001 |
| *oqx**B* | Decrease | < 0.001 | < 0.001 |
| *Arn**T* | Decrease | < 0.001 | < 0.001 |
| *oqx**A* | Decrease | < 0.001 | < 0.001 |
| *ept**B* | Decrease | < 0.001 | < 0.001 |
| *Lpt**D* | Decrease | < 0.001 | < 0.001 |
| *tet*(*Q*) | Increase | < 0.001 | < 0.001 |
| *APH*(3')-*Ia* | Decrease | < 0.001 | < 0.001 |
| *Klebsiella pneumoniae*<br>*Kpn**G* | Decrease | < 0.001 | < 0.001 |
| *Omp**A* | Decrease | < 0.001 | < 0.001 |
| *OXY*-*I*-3 | Decrease | < 0.001 | < 0.001 |
| *Fos**A6* | Decrease | < 0.001 | < 0.001 |
| *OXY*-*I*-4 | Decrease | < 0.001 | < 0.001 |
| *Klebsiella pneumoniae*<br>*Kpn**F* | Decrease | < 0.001 | < 0.001 |
| *Klebsiella pneumoniae*<br>*Kpn**E* | Decrease | 0.001 | 0.006 |
| *bac**A* | Decrease | 0.001 | 0.006 |
| *Klebsiella pneumoniae*<br>*Omp**K37* | Decrease | 0.003 | 0.015 |
| *H*-*NS* | Decrease | 0.007 | 0.029 |
| *emr**K* | Increase | 0.007 | 0.030 |
| *mdt**O* | Increase | 0.008 | 0.030 |
| *mdt**A* | Increase | 0.009 | 0.034 |
| *mdt**N* | Increase | 0.010 | 0.037 |
| *gad**W* | Increase | 0.012 | 0.043 |
| *mdt**C* | Decrease | 0.013 | 0.044 |
| *Acr**S* | Increase | 0.013 | 0.044 |
| *Escherichia coli* *mdf**A* | Increase | 0.018 | 0.057 |
| *CRP* | Decrease | 0.020 | 0.058 |
| *vanS* gene in *vanD* cluster | Increase | 0.019 | 0.058 |
| *evg**A* | Increase | 0.020 | 0.058 |
| *mdt**F* | Increase | 0.021 | 0.061 |
| *emr**Y* | Increase | 0.025 | 0.067 |
| *leu**O* | Increase | 0.027 | 0.071 |
| *mdt**E* | Increase | 0.035 | 0.089 |
| *Cep**A*-49 | Increase | 0.040 | 0.091 |
| *emr**R* | Increase | 0.040 | 0.091 |
| *Pmr**F* | Increase | 0.049 | 0.108 |
| *Male* |  |  |  |
| *APH*(3')-*Ia* | Decrease | < 0.001 | < 0.001 |
| *Lpt**D* | Decrease | < 0.001 | < 0.001 |
| *tet*(*Q*) | Increase | < 0.001 | < 0.001 |
| *Arn**T* | Decrease | < 0.001 | < 0.001 |
| *ept**B* | Decrease | < 0.001 | < 0.001 |
| *Klebsiella pneumoniae*<br>*Kpn**G* | Decrease | < 0.001 | < 0.001 |
| *oqxB* | Decrease | < 0.001 | < 0.001 |
| *OXY-1-4* | Decrease | < 0.001 | < 0.001 |
| *Klebsiella pneumoniae*<br>*acrA* | Decrease | < 0.001 | < 0.001 |
| *bacA* | Decrease | < 0.001 | < 0.001 |
| *OmpA* | Decrease | < 0.001 | < 0.001 |
| *Klebsiella pneumoniae*<br>*KpnF* | Decrease | < 0.001 | < 0.001 |
| *FosA6* | Decrease | < 0.001 | < 0.001 |
| *Klebsiella pneumoniae*<br>*KpnE* | Decrease | < 0.001 | < 0.001 |
| *oqxA* | Decrease | < 0.001 | < 0.001 |
| *vanR* gene in *vanD* cluster | Increase | < 0.001 | < 0.001 |
| *Klebsiella pneumoniae*<br>*KpnH* | Decrease | < 0.001 | < 0.001 |
| *Klebsiella pneumoniae*<br>*OmpK37* | Decrease | < 0.001 | 0.001 |
| *vanS* gene in *vanD* cluster | Increase | < 0.001 | 0.001 |
| *emrA* | Decrease | < 0.001 | 0.002 |
| *OXY-1-3* | Decrease | < 0.001 | 0.002 |
| *tet(W)* | Increase | 0.002 | 0.006 |
| *tet(W/N/W)* | Increase | 0.004 | 0.012 |
| *vanH* gene in *vanD* cluster | Increase | 0.008 | 0.024 |
| *vanV* gene in *vanB* cluster | Increase | 0.010 | 0.027 |
| *H-NS* | Decrease | 0.022 | 0.058 |
| *mdtC* | Decrease | 0.033 | 0.081 |
| *TolC* | Increase | 0.039 | 0.095 |
| *msrC* | Increase | 0.050 | 0.117 |

**Table S4.** ARGs with an abundance greater than 1% that significantly change with time (1-22 months) in the Unaltered group. No genera changed significantly over time in the Unaltered female mice.

| ARG | Change over time | p-value | q-value |
| --- | --- | --- | --- |
| *Female* |  |  |  |
| *tet(O/32/O)* | Increase | 0.006 | 0.159 |
| *tet(W/N/W)* | Increase | 0.006 | 0.159 |
| *Male* |  |  |  |
| *tet(40)* | Increase | 0.002 | 0.100 |

**Table S5.**
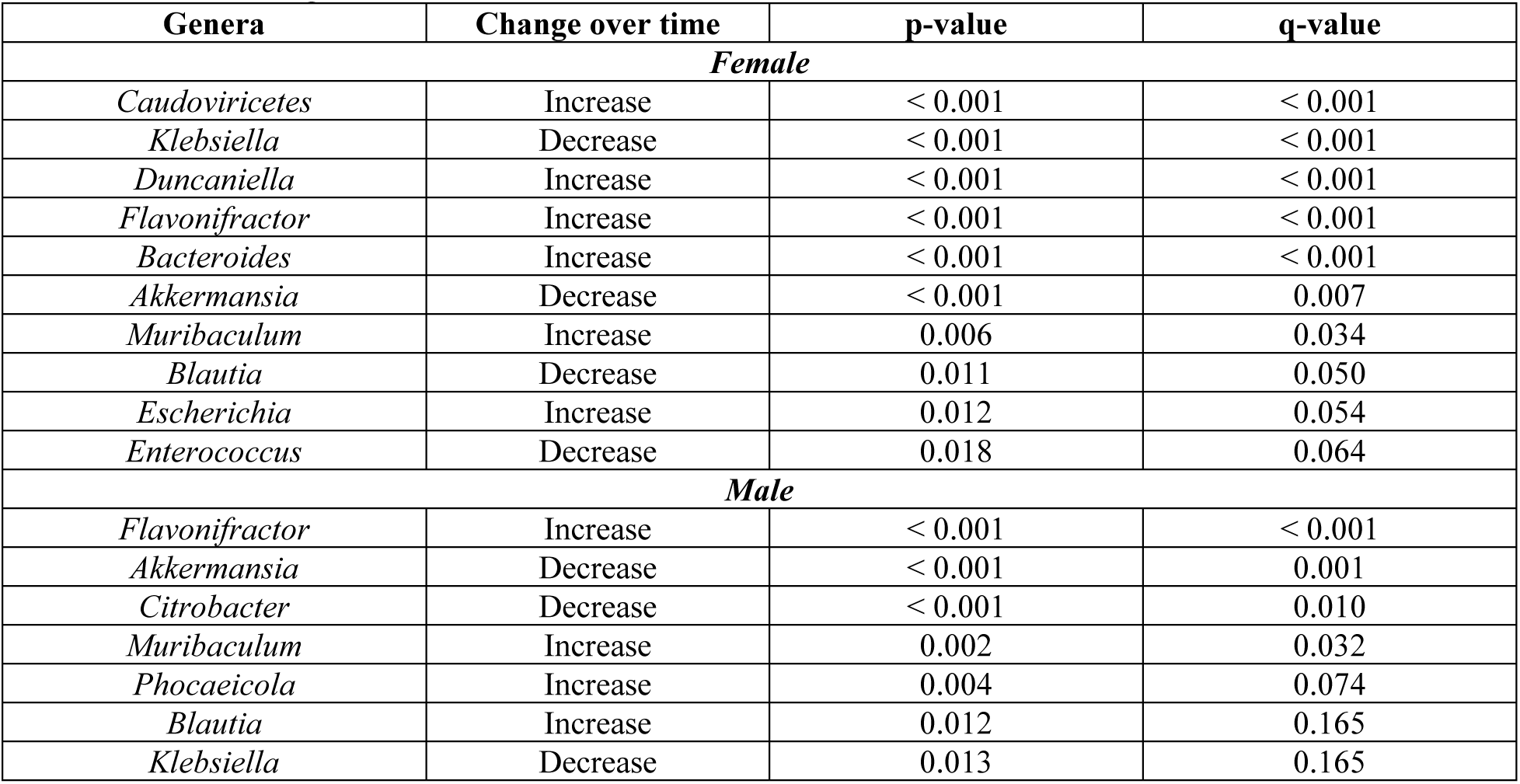
Genera that significantly changed with time (3-22 months) in the Continuous Dose group with a relative abundance greater than 1%.

**Table S6.** Genera that significantly changed with time (1-22 months) in the Unaltered group with an abundance greater than 1%. Unaltered female mice did not have any genera that significantly changed with time.

| Genera | Change over time | p-value | q-value |
| --- | --- | --- | --- |
| *Male* |  |  |  |
| *Muribaculaceae bacterium* | Decrease | < 0.001 | < 0.001 |
| *Faecalibaculum* | Decrease | 0.001 | 0.056 |
| *[Clostridium]* | Increase | 0.019 | 0.165 |
| *Blautia* | Increase | 0.018 | 0.165 |
| *Caudoviricetes* | Increase | 0.019 | 0.165 |
| *Muribaculum* | Decrease | 0.017 | 0.165 |
| *Lachnoclostridium* | Increase | 0.028 | 0.185 |
| *Roseburia* | Increase | 0.030 | 0.190 |
| *Acutalibacter* | Increase | 0.035 | 0.204 |
| *Enterocloster* | Increase | 0.038 | 0.208 |

**Table S7.** Variance in the % of ARG abundance associated with *Enterobacteriaceae* family amongst treatment groups over time. Shapiro’s test was used to check for the normality of each dataset. A Likelihood Ratio Test was used comparing two Generalized Least Squares models with homoscedastic and heteroscedastic residual variance to determine changes in variance over time.

| Sex | Group | Age (Months) |  |  |  |  |  |  |  | p value |
| --- | --- | --- | --- | --- | --- | --- | --- | --- | --- | --- |
|  |  | 1 | 3 | 5 | 13 | 18 | 18.5 | 20 | 22 |  |
| Female | Unaltered |  | 7.044 | 398.242 | 0.821 | 81.225 |  |  | 99.571 | <0.001 |
|  | Continuous Dose | 550.624 | 10.871 | 1122.147 | 31.780 | 296.521 |  |  | 311.983 | 0.003 |
|  | Delayed Dose |  | 55.596 | 814.657 | 4.248 | 20.991 | 3.127 | 457.193 | 159.998 | <0.001 |
|  | Initial Dose | 27.876 | 14.713 | 392.422 | 83.987 | 368.685 | 63.367 | 42.202 | 43.186 | <0.001 |
| Male | Unaltered |  | 94.663 | 46.378 | 132.576 | 63.907 |  |  | 54.000 | 0.866 |
|  | Continuous Dose | 1.342 | 13.083 | 60.052 | 31.834 | 47.476 |  |  | 65.539 | 0.385 |
|  | Delayed Dose |  | 14.805 | 72.633 | 1.182 | 337.242 | 44.286 | 275.398 | 297.194 | <0.001 |
|  | Initial Dose | 2.233 | 6.099 | 49.266 | 59.217 | 213.520 | 59.571 | 129.980 | 447.772 | <0.001 |

**Table S8.** Variance in the ARG richness and abundance amongst treatment groups over time. Shapiro’s test was used to check for the normality of each dataset. A Likelihood Ratio Test was used comparing two Generalized Least Squares models with homoscedastic and heteroscedastic residual variance to determine changes in variance over time.

| Sex | Group | Age (Months) |  |  |  |  |  |  |  | p value |
| --- | --- | --- | --- | --- | --- | --- | --- | --- | --- | --- |
|  |  | 1 | 3 | 5 | 13 | 18 | 18.5 | 20 | 22 |  |
| **ARG richness** |  |  |  |  |  |  |  |  |  |  |
| Female | Unaltered |  | 1.667 | 365.667 | 6.333 | 14.054 |  |  | 32.691 | < 0.001 |
|  | Continuous Dose | 112.500 | 24.333 | 2.333 | 134.250 | 81.200 |  |  | 74.992 | 0.213 |
|  | Delayed Dose |  | 110.250 | 426.000 | 28.667 | 37.242 | 14.167 | 180.515 | 222.981 | 0.004 |
|  | Initial Dose | 4.500 | 5.667 | 489.667 | 7.583 | 199.788 | 58.447 | 55.242 | 12.692 | < 0.001 |
| Male | Unaltered |  | 40.000 | 21.583 | 72.333 | 42.233 |  |  | 30.374 | 0.867 |
|  | Continuous Dose | 4.500 | 4.250 | 6.250 | 15.000 | 26.386 |  |  | 26.218 | 0.369 |
|  | Delayed Dose |  | 9.000 | 13.667 | 4.917 | 268.691 | 97.200 | 121.356 | 165.474 | 0.006 |
|  | Initial Dose | 0.500 | 8.250 | 18.000 | 4.000 | 22.164 | 51.818 | 106.750 | 63.381 | 0.007 |
| **ARG abundance** |  |  |  |  |  |  |  |  |  |  |
| Female | Unaltered |  | 4.102 | 74.315 | 6.383 | 11.449 |  |  | 3.484 | 0.002 |
|  | Continuous Dose | 13.468 | 750.200 | 41.709 | 77.982 | 93.890 |  |  | 58.519 | 0.011 |
|  | Delayed Dose |  | 3.740 | 45.690 | 13.091 | 11.236 | 1118.776 | 1378.560 | 1946.852 | < 0.001 |
|  | Initial Dose | 5.848 | 128.480 | 83.415 | 9.152 | 105.099 | 11.877 | 7.222 | 10.566 | < 0.001 |
| Male | Unaltered |  | 2.026 | 2.735 | 0.417 | 21.294 |  |  | 20.488 | 0.014 |
|  | Continuous Dose | 0.120 | 78.237 | 10.136 | 19.831 | 12.289 |  |  | 23.834 | 0.065 |
|  | Delayed Dose |  | 4.927 | 12.675 | 4.615 | 45.944 | 1805.357 | 246.861 | 195.395 | < 0.001 |
|  | Initial Dose | 31.920 | 103.841 | 13.030 | 0.337 | 29.421 | 4.299 | 6.183 | 221.139 | < 0.001 |

**Table S9.**
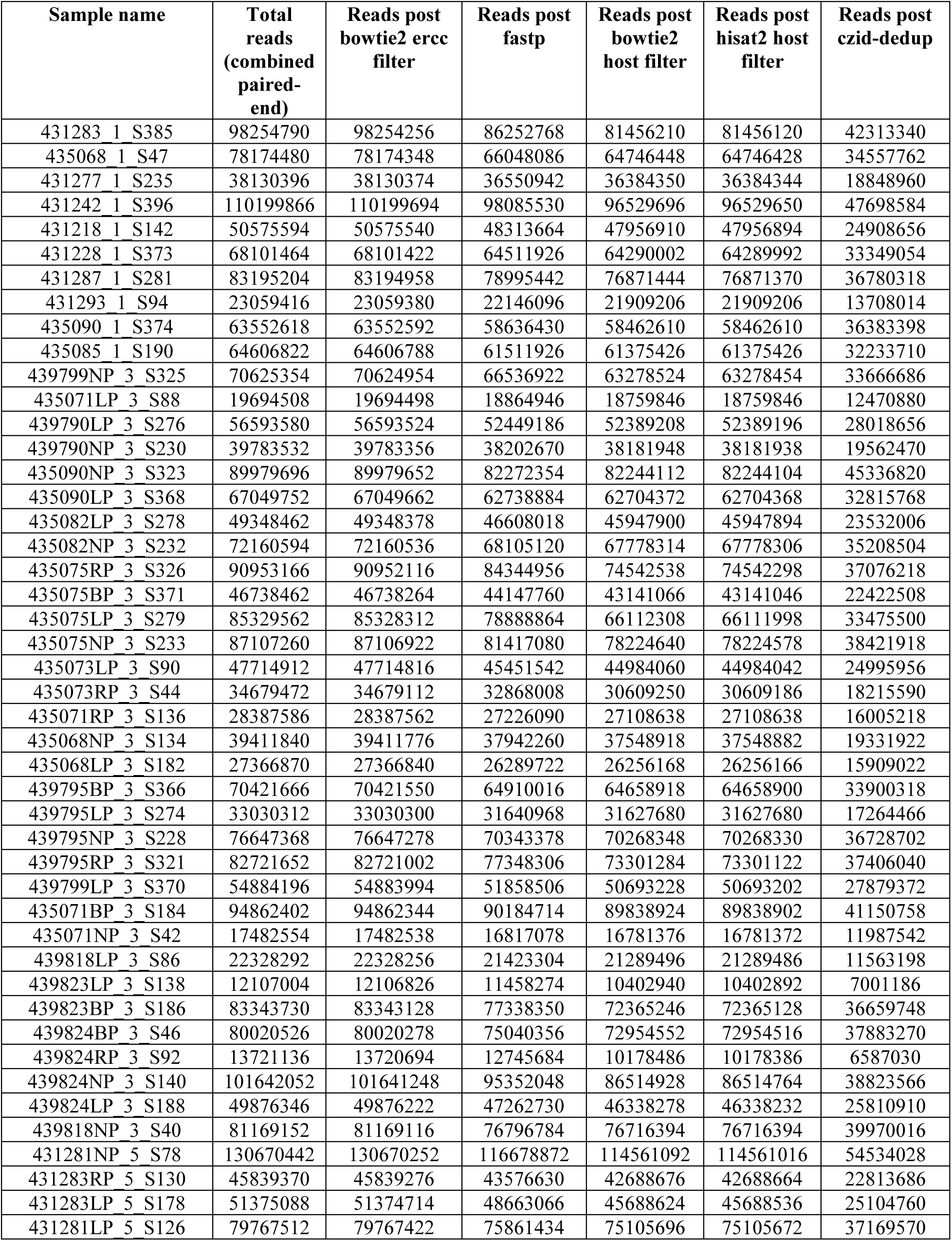
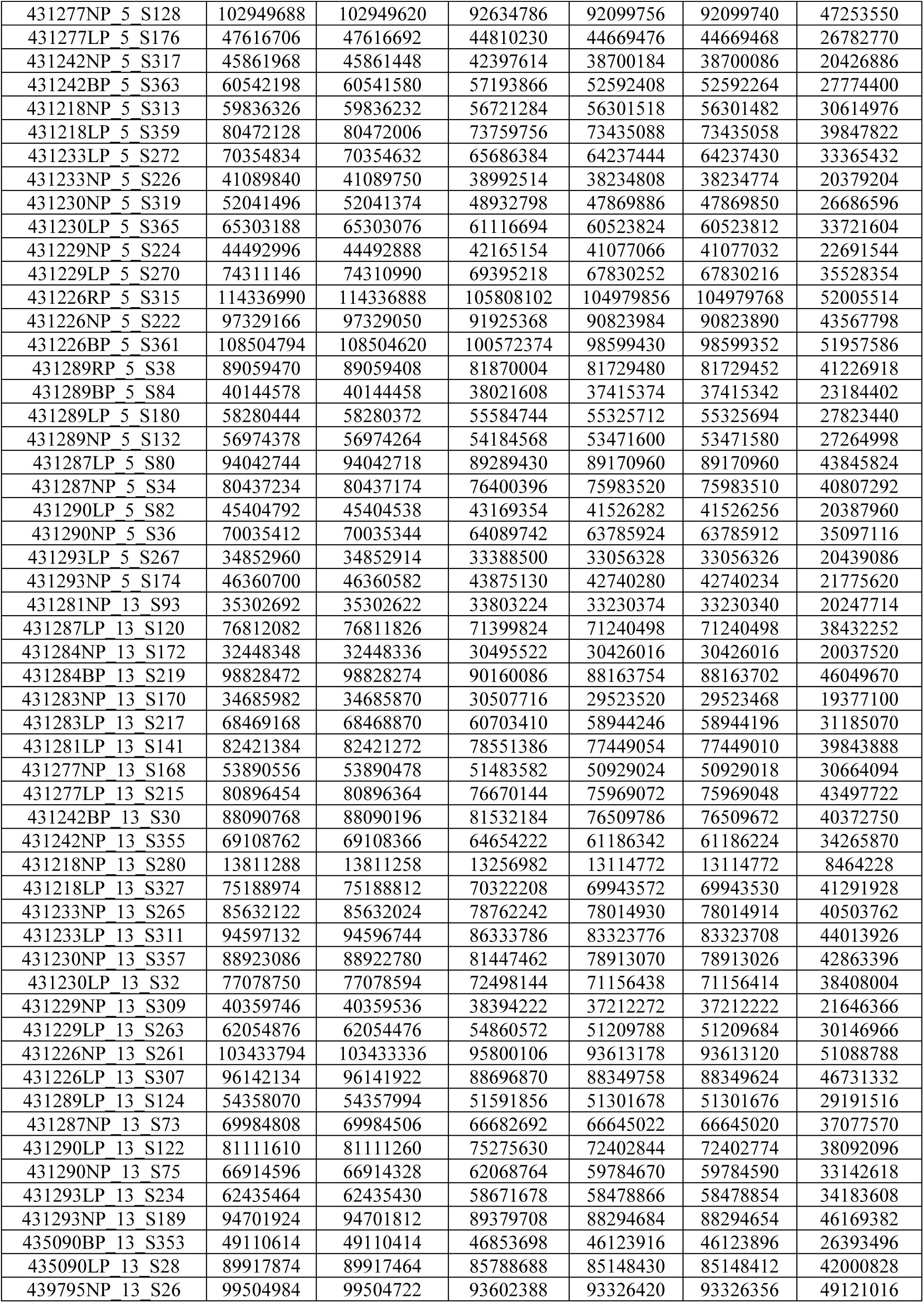
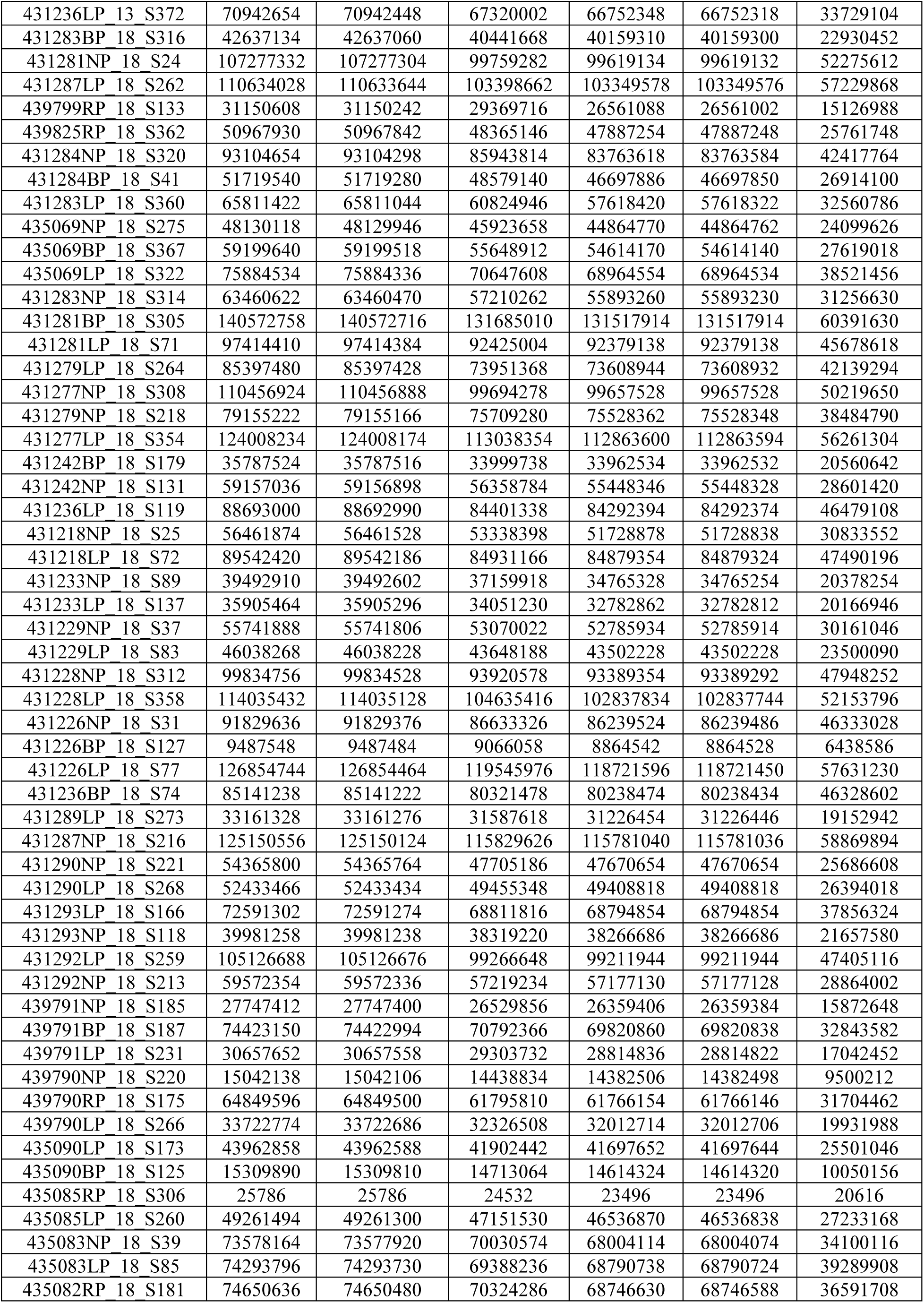
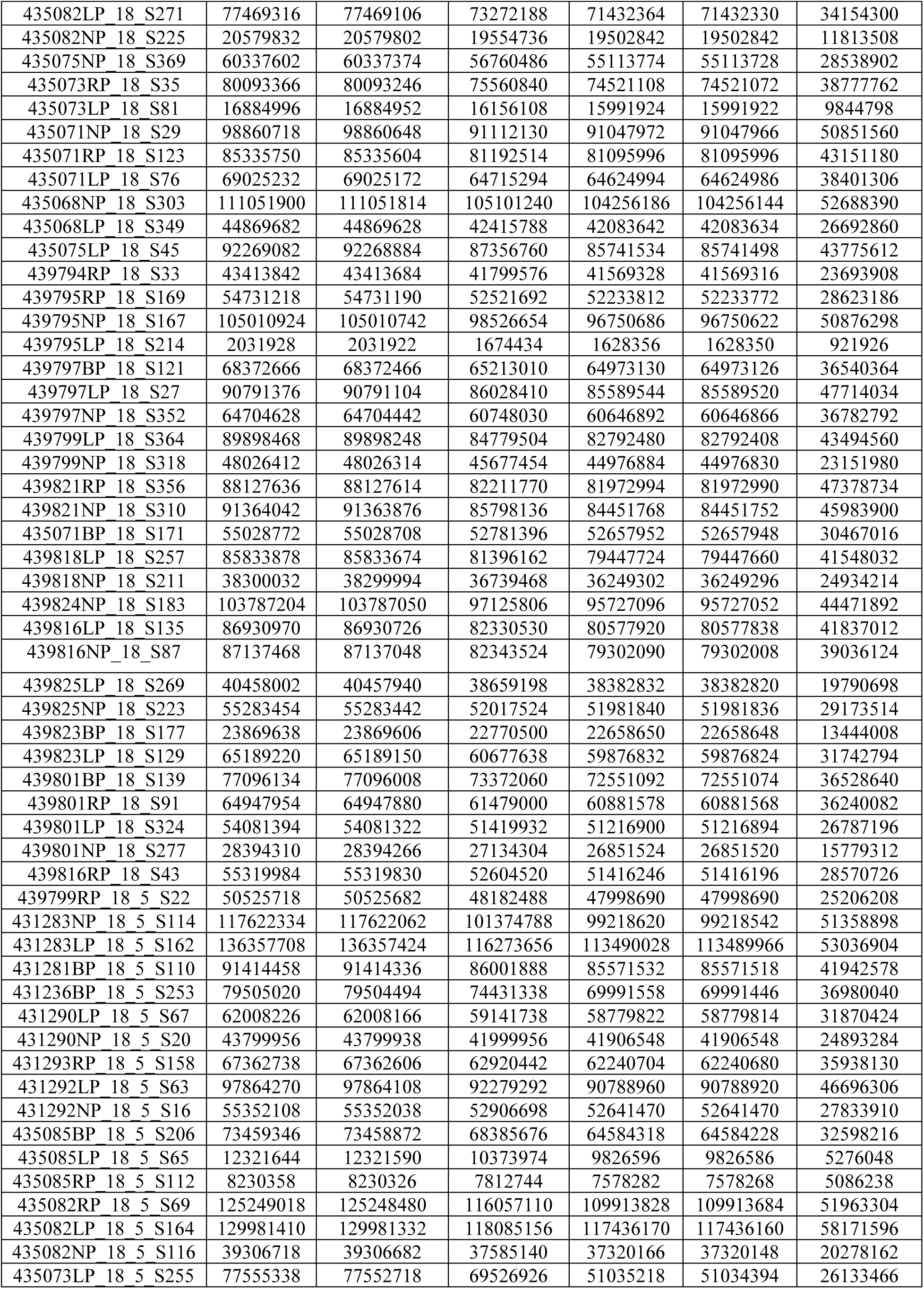
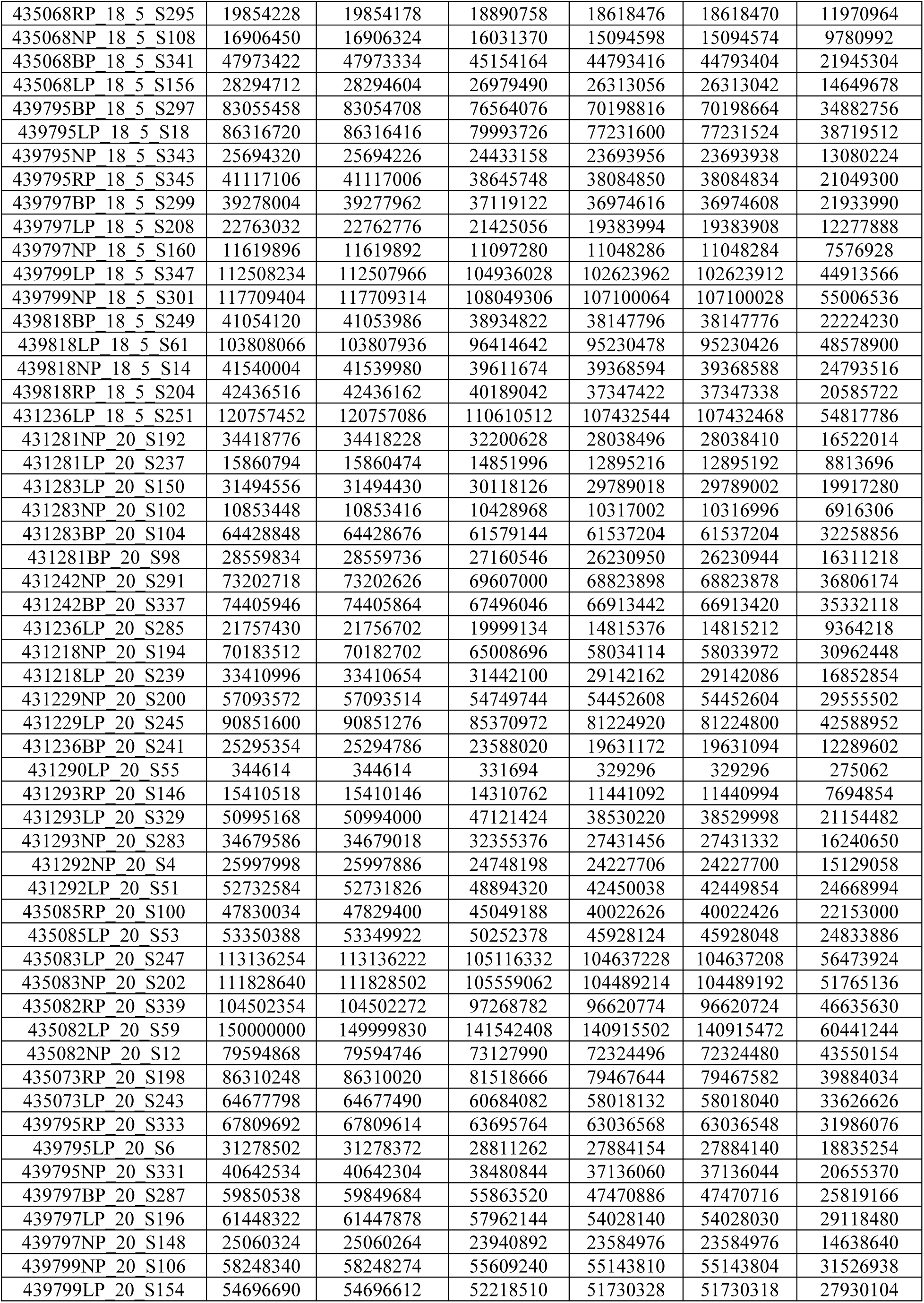
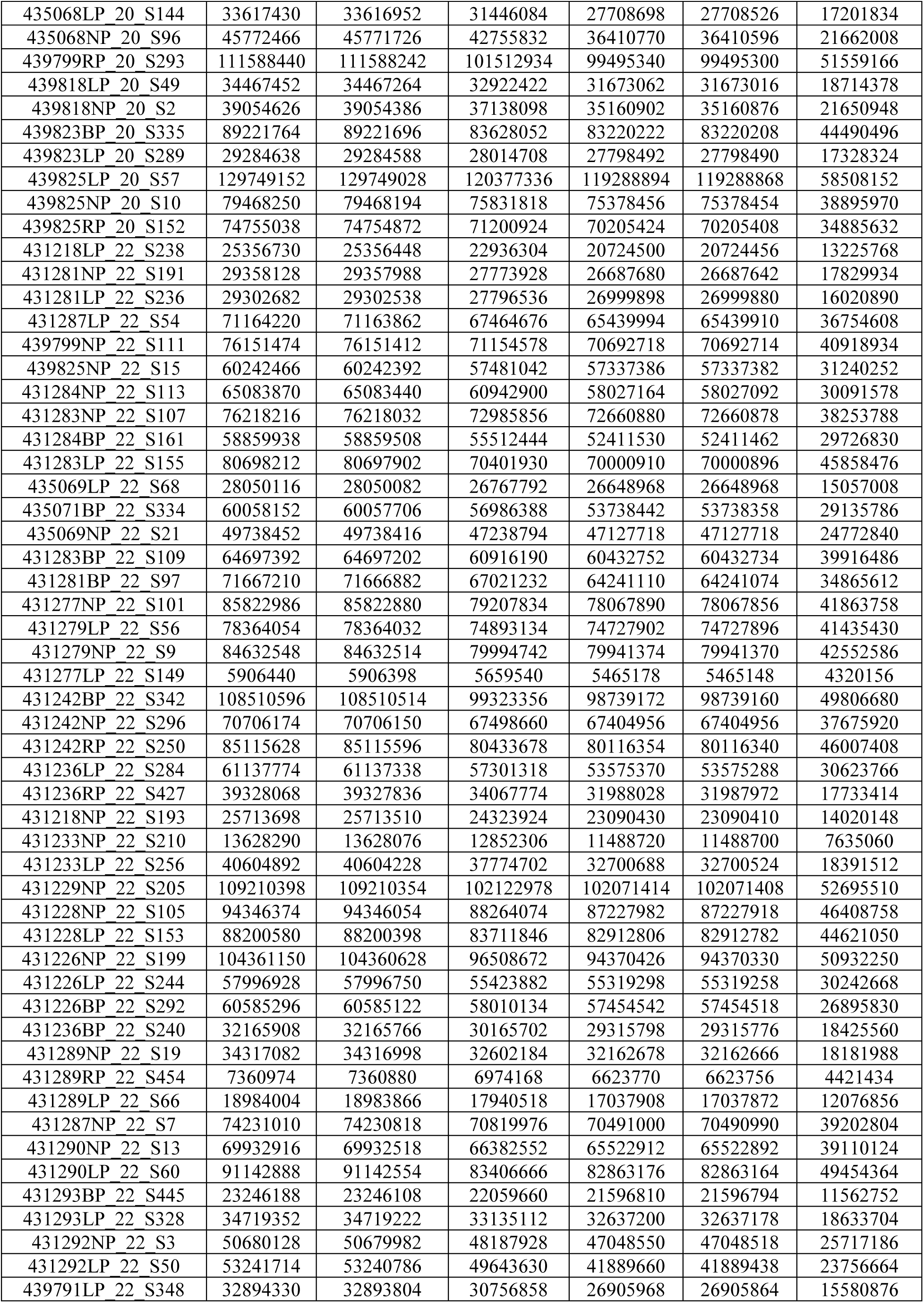
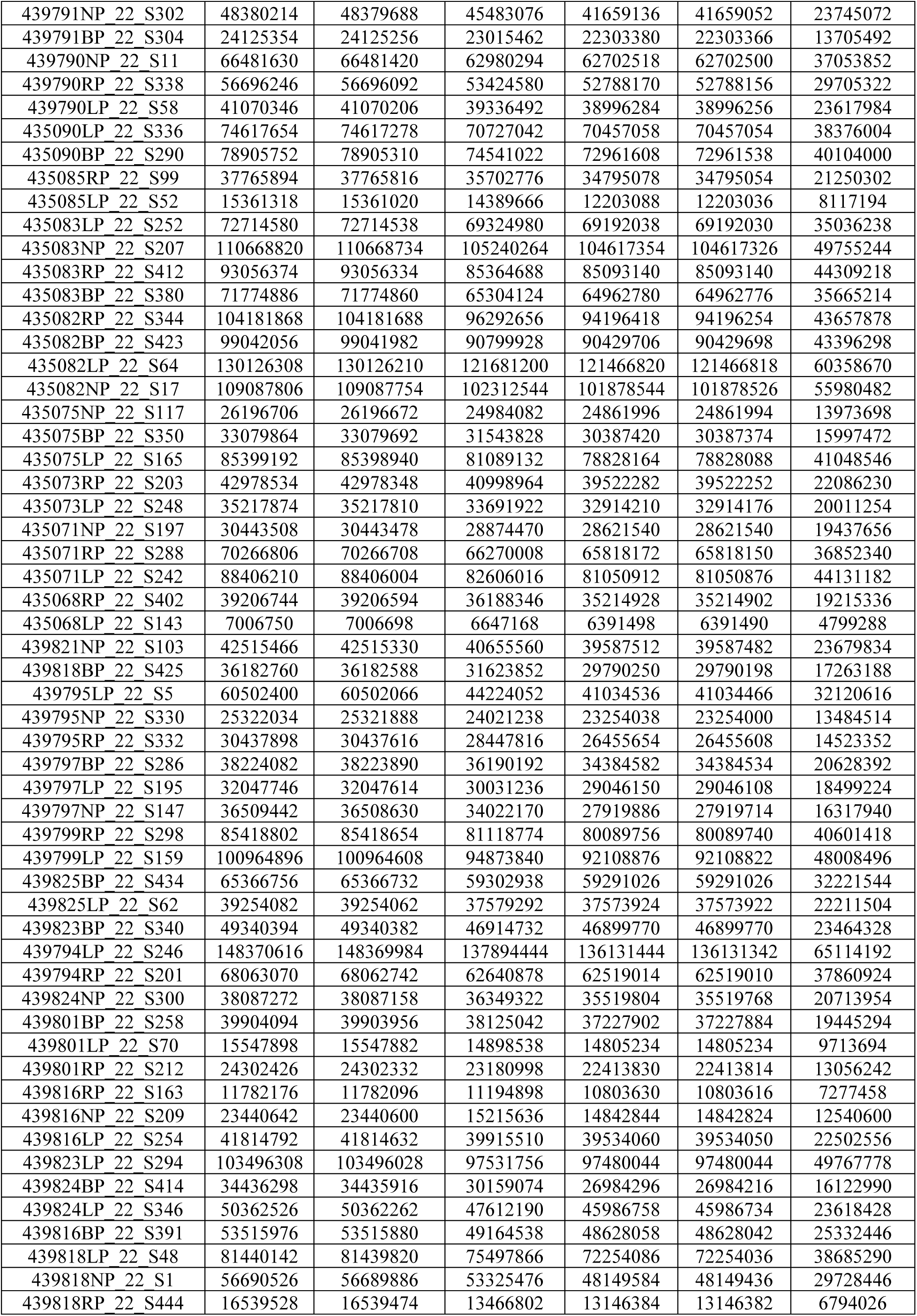
Quality control processing of shotgun metagenomic paired-end samples using the Chan Zuckerberg ID pipeline.

**Table S10.**
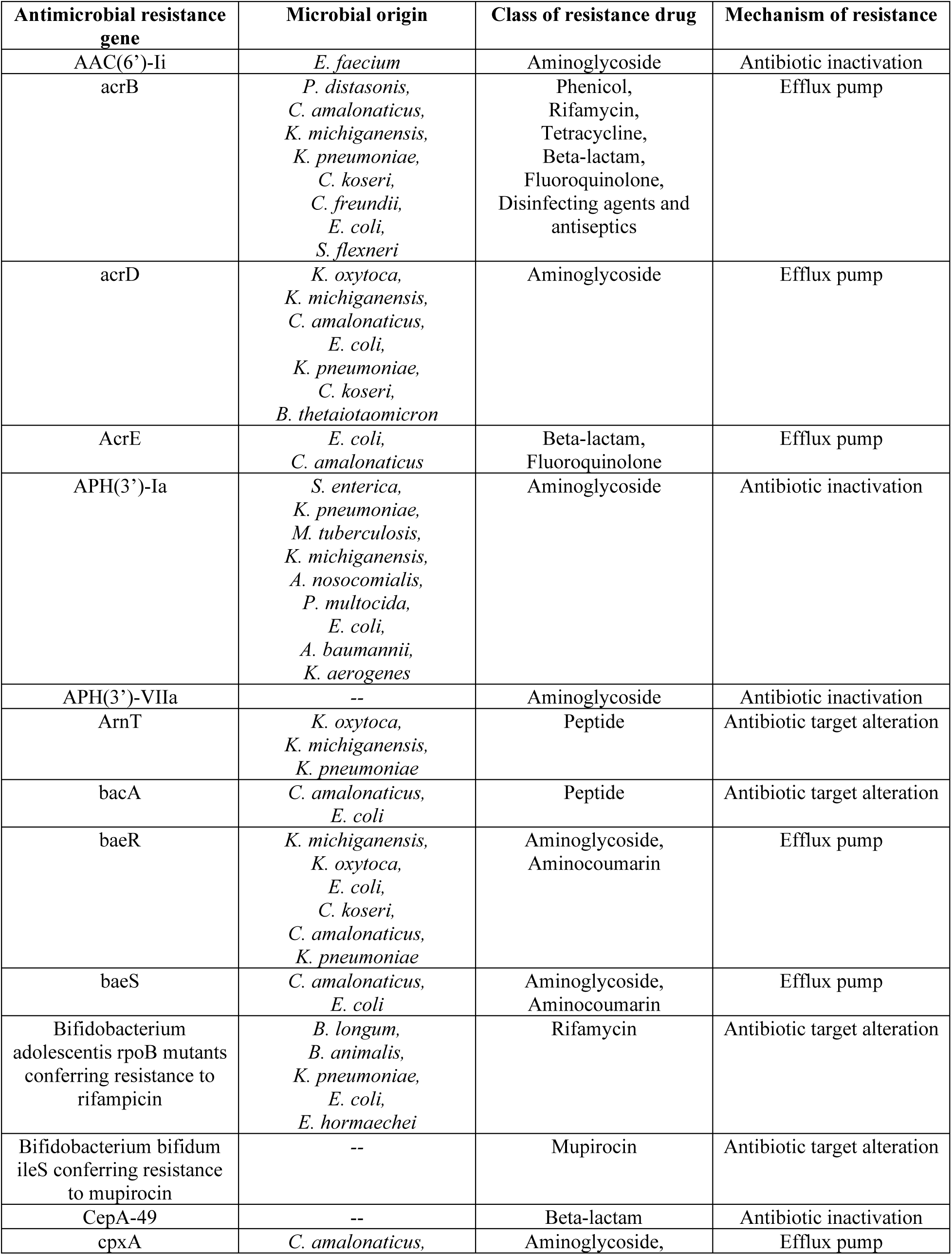
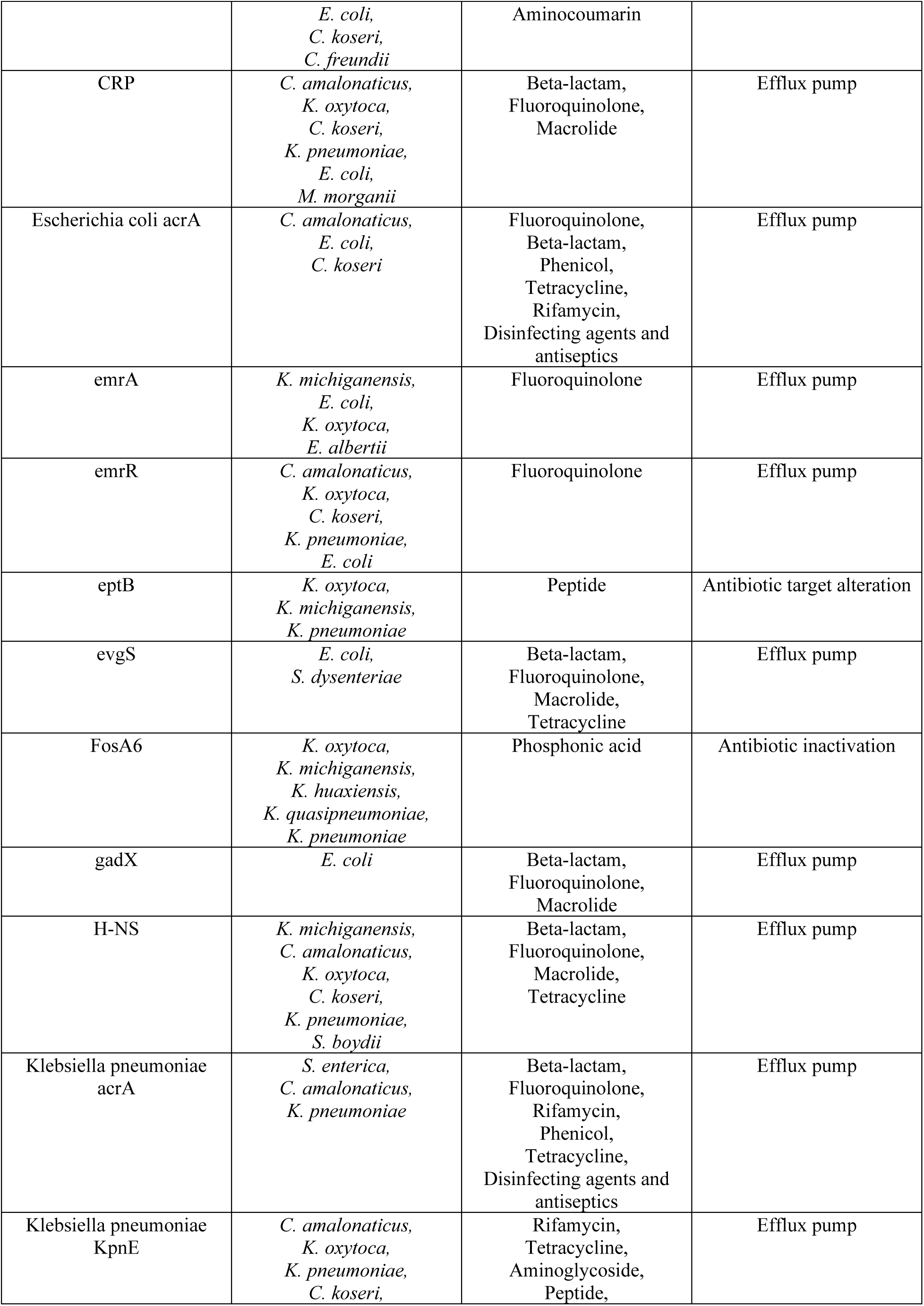
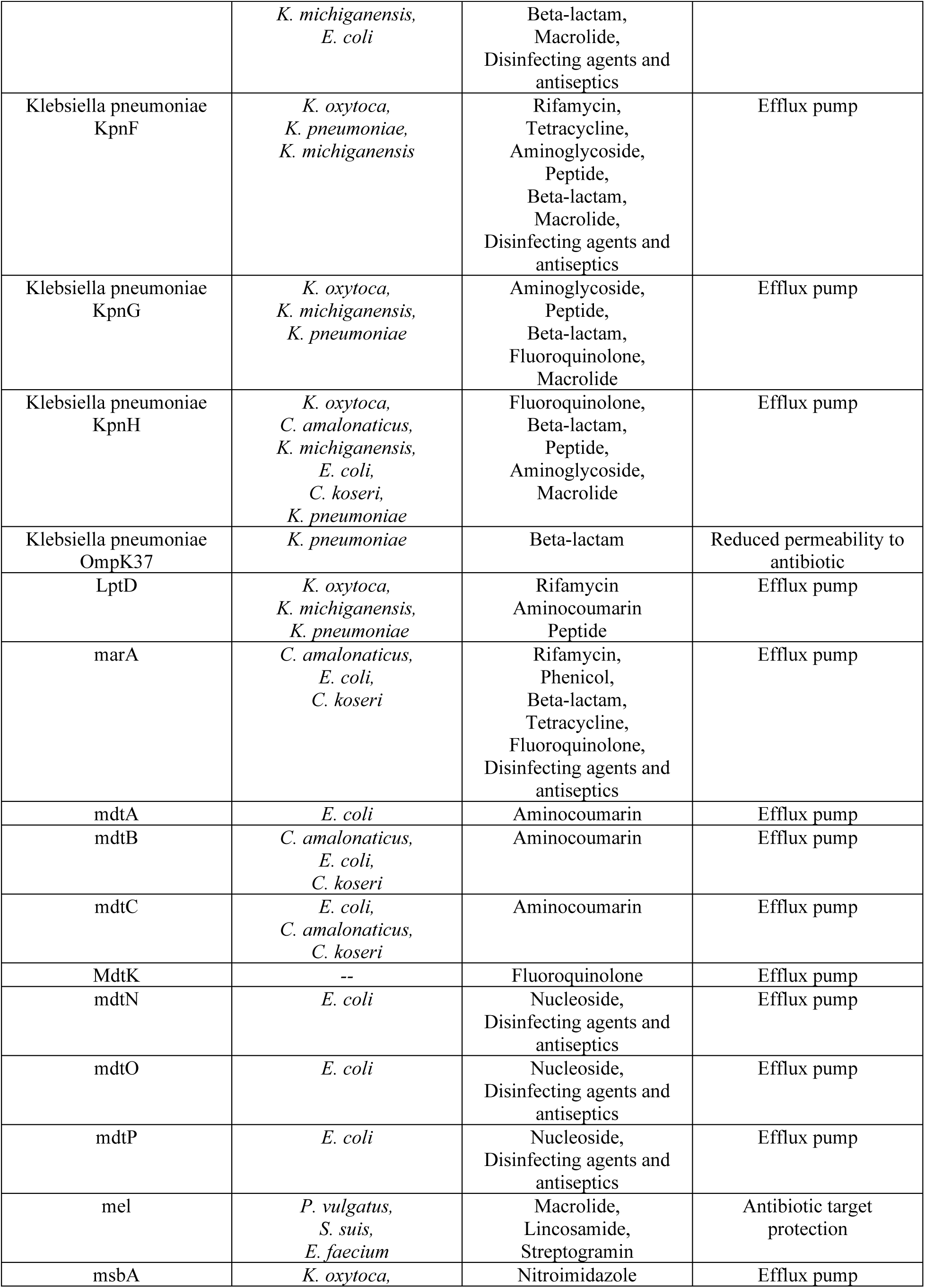
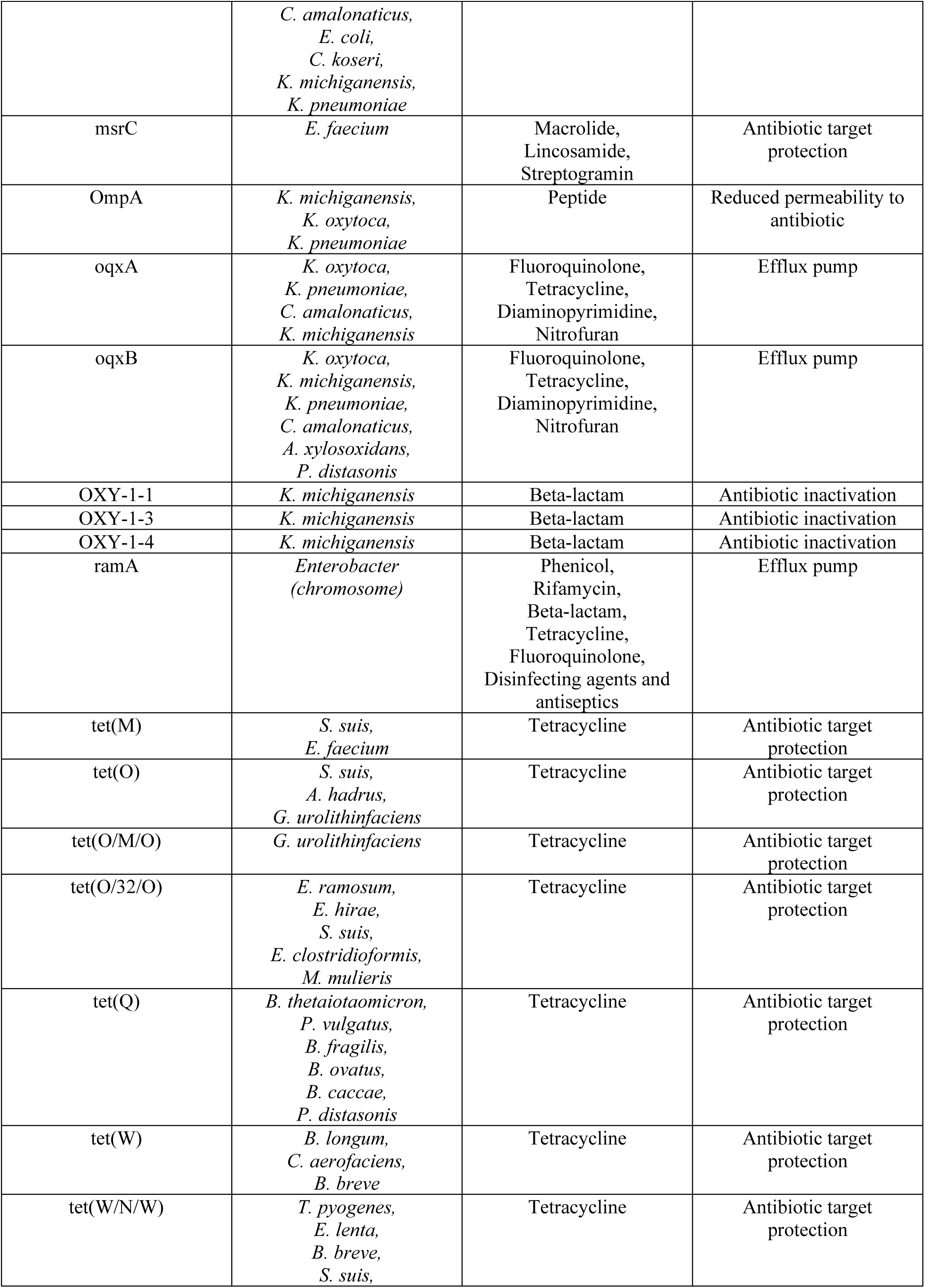
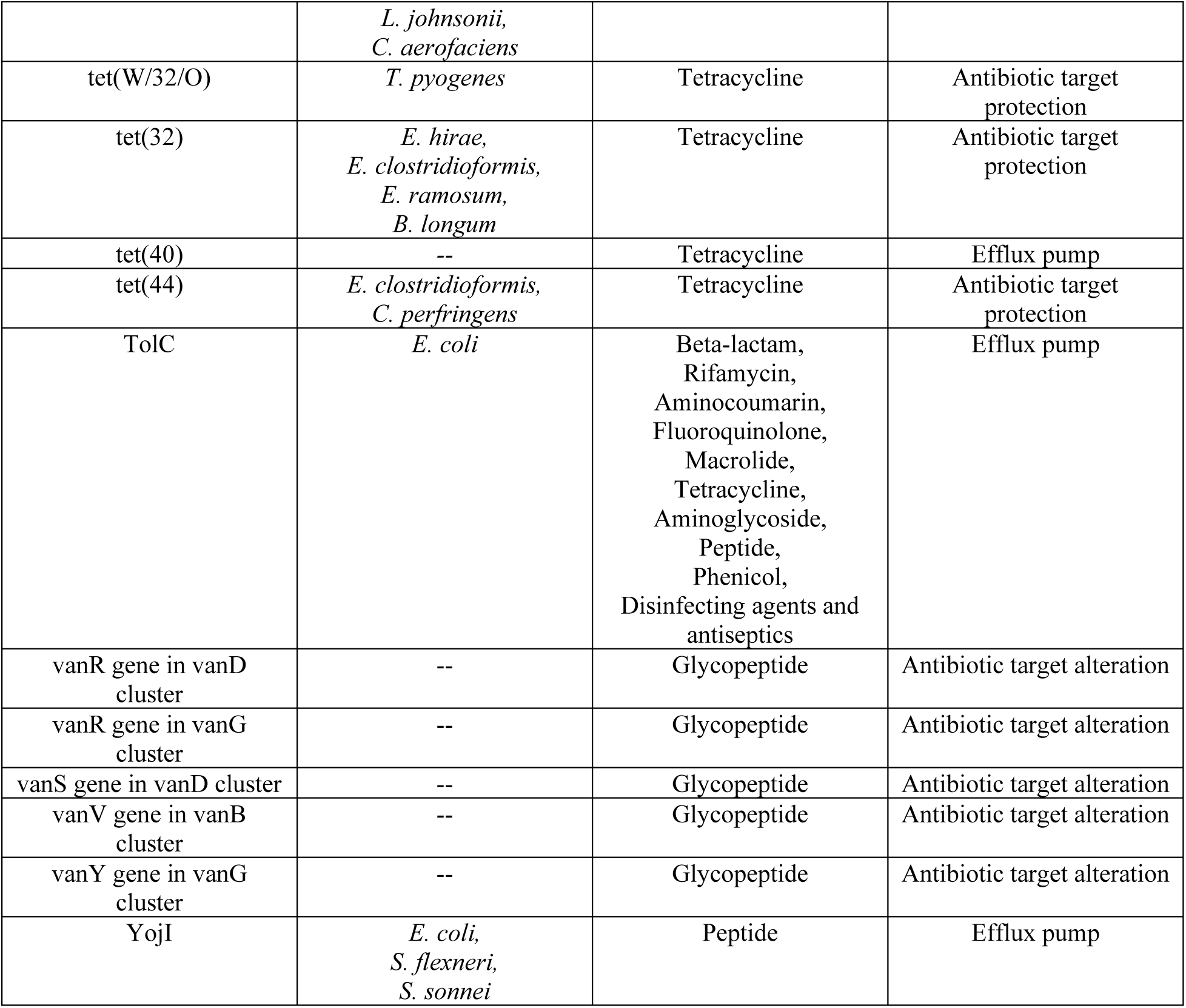
Microbial origin and classification of antimicrobial resistance genes (> 1% relative abundance) detected in female and male mice in overall study.

## References and Notes

1. Antimicrobial Resistance Collaborators, Global burden of bacterial antimicrobial resistance in 2019: a systematic analysis. Lancet 399, 629–655 (2022). doi: 10.1016/S0140-6736(21)02724-0.

2. V.T. Chu, A. Glascock, D. Donnell, C. Grabow, C.E. Brown, R. Ward, C. Love, K.L. Kalantar, S.E. Cohen, C. Cannon, M.H. Woodworth, C.F. Kelley, C. Celum, A.F Leutkemeyer, C.R. Langelier, Impact of doxycycline post-exposure prophylaxis for sexually transmitted infections on the gut microbiome and antimicrobial resistome. Nat. Med. 31, 2017–217 (2025). doi: 10.1038/s41591-024-03274-2.

3. R.S. McInnes, G.E. McCallum, L.E. Lamberte, W. van Schaik, Horizontal transfer of antibiotic resistance genes in the human gut microbiome. Curr. Opin. Microbiol. 53, 35–43 (2020). doi: 10.1016/j.mib.2020.02.002.

4. D.G.J. Larsson, C.F. Flach, Antibiotic resistance in the environment. Nat. Rev. Microbiol. 20, 257–269 (2022). doi: 10.1038/s41579-021-00649-x.

5. I.L. Brito, S. Yilmaz, K. Huang, L. Xu, S.D. Jupiter, A.P. Jenkins, W. Naisilisili, M. Tamminen, C.S. Smillie, J.R. Wortman, B.W. Birren, R.J. Xavier, P.C. Blainey, A.K Singh, D. Gevers, E.J. Alm, Mobile genes in the human microbiome are structured from global to individual scales. Nature 535, 435–439 (2016). doi: 10.1038/nature18927.

6. P.J. Diebold, M.W. Rhee, Q. Shi, N.V. Trung, F. Umrani, S. Ahmed, V. Kulkarni, P. Deshpande, M. Alexander, N.T. Hoa, N.A. Christakis, N.T. Iqbal, S.A. Ali, J.S. Mathad, I.L. Brito, Clinically relevant antibiotic resistance genes are linked to a limited set of taxa within gut microbiome worldwide. Nat. Comm. 14, 7366 (2023). doi: 10.1038/s41467-023-42998-6.

7. E.C. Pehrsson, P. Tsukayama, S. Patel, M. Mejía-Bautista, G. Sosa-Soto, K.M. Navarrete, M. Calderon, L. Cabrera, W. Hoyos-Arango, M.T. Bertoli, D.E. Berg, R.H. Gilman, G. Dantas, Interconnected microbiomes and resistomes in low-income human habitats. Nature 533, 212–216 (2016). doi: 10.1038/nature17672.

8. A. Palleja, K.H. Mikkelsen, S.K Forslund, A. Kashani, K.H. Allin, T. Nielsen, T.H. Hansen, S. Liang, Q. Feng, C. Zhang, P.T. Pyl, L.P. Coelho, H. Yang, J. Wang, A. Typas, M.F. Nielsen, H.B. Nielsen, P. Bork, J. Wang, T. Vilsbøll, T. Hansen, F.K. Knop, M. Arumugam, O. Pedersen, Recovery of gut microbiota of healthy adults following antibiotic exposure. Nat. Microbiol. 3, 1255–1265 (2018). doi: 10.1038/s41564-018-0257-9.

9. A. Rashidi, M. Ebadi, T.U. Rehman, H. Elhusseini, D. Kazadi, H. Halaweish, M.H. Khan, A. Hoeschen, Q. Cao, X. Luo, A.J. Kabage, S. Lopez, S.G. Holtan, D.J. Weisdorf, C. Liu, S. Ishii, A. Khoruts, C. Staley, Long- and short-term effects of fecal microbiota transplantation on antibiotic resistance genes: results from a randomized placebo-controlled trial. Gut Microbes 16, 2327442 (2024). doi: 10.1080/19490976.2024.2327442.

10. K. Lee, S. Raguideau, K. Sirén, F Asnicar, F. Cumbo, F. Hildebrand, N. Segata, C.J. Cha, C. Quince, Population-level impacts of antibiotic usage on the human gut microbiome. Nat. Comm. 14, 1191 (2023). doi: 10.1038/s41467-023-36633-7.

11. A.M. Turner, L. Li, I.R. Monk, J.Y.H. Lee, D.J. Ingle, S. Portelli, N.L. Sherry, N. Isles, T. Seemann, L.K. Sharkey, C.J. Walsh, G.E Reid, S. Nie, B.A. Eijkelkamp, N.E. Holmes, B. Collis, S. Vogrin, A. Hiergeist, D. Weber, A. Gessner, E. Holler, D.B. Ascher, S. Duchene, N.E. Scott, T.P. Stinear, J.C. Kwong, C.L. Gorrie, B.P. Howden, G.P. Carter, Rifaximin prophylaxis causes resistance to the last-resort antibiotic daptomycin. Nature 635, 969–977 (2024). doi: 10.1038/s41586-024-08095-4.

12. K.S. Xue, S.J. Walton, D.A. Goldman, M.L. Morrison, A.J. Verster, A.B. Parrott, F.B. Yu, N.F. Neff, N.A. Rosenberg, B.D. Ross, D.A. Petrov, K.C. Huang, B.H. Good, D.A. Relman, Prolonged delays in human microbiota transmission after a controlled antibiotic perturbation. *bioRxiv* 2023.09.26.559480 (2023). doi: 10.1101/2023.09.26.559480.

13. J. Suez, N. Zmora, G. Zilberman-Schapira, U. Mor, M. Dori-Bachash, S. Bashiardes, M. Zur, D. Regev-Lehavi, R.B.Z. Brik, S. Federici, M. Horn, Y. Cohen, A.E. Moor, D. Zeevi, T. Korem, E. Kotler, A. Harmelin, S. Itzkovitz, N. Maharshak, O. Shibolet, M. Pevsner-Fischer, H. Shapiro, I. Sharon, Z. Halpern, E. Segal, E. Elinav, Post-antibiotic gut mucosal microbiome reconstitution is impaired by probiotics and improved by autologous FMT. Cell 174, 1406–1423 (2018). doi: 10.1016/j.cell.2018.08.047.

14. A.J. Gasparrini, B. Wang, X. Sun, E.A. Kennedy, A. Hernandez-Leyva, I.M. Ndao, P.I. Tarr, B.B. Warner, G. Dantas, Metagenomic signatures of early life hospitalization and antibiotic treatment in the infant gut microbiota and resistome persist long after discharge. Nat. Microbiol. 4, 2285–2297 (2019). doi: 10.1038/s41564-019-0550-2.

15. J.S.Y. Lau, T.M. Korman, I. Woolley, Life-long antimicrobial therapy: where is the evidence? J. Antimicrob. Chemother. 73, 2601–2612 (2018). doi: 10.1093/jac.dky174.

16. T.R. Weiner, D.B. El-Najjar, C.L. Herndon, C.C. Wyles, H.J. Cooper, How are oral antibiotics being used in total joint arthroplasty? A review of the literature. Orthop. Rev. 16, 92287 (2024). doi: 10.52965/001c.92287.

17. K.S. O’Brien, A.M. Arzika, A. Amza, R. Maliki, B. Aichatou, I.M. Bello, D. Beidi, N. Galo, N. Harouna, A.M. Karamba, S. Mahamadou, M. Abarchi, A. Ibrahim, E. Lebas, B. Peterson, Z. Liu, V. Le, E. Colby, T. Doan, J.D. Keenan, C.E. Oldenburg, T.C. Porco, B.F. Arnold, T.M. Lietman, AVENIR study group, Azithromycin to reduce mortality – an adaptive cluster-randomized trial. N. Engl. J. Med. 391, 699–709 (2024). doi: 10.1056/NEJMoa2312093.

18. S.K. Bhattarai, M. Du, A.L. Zeamer, B.M. Morzfeld, T.D. Kellogg, K. Firat, A. Benjamin, J.M. Bean, M. Zimmerman, G. Mardi, S.C. Vilbrun, K.F. Walsh, D.W. Fitzgerald, M.S. Glickman, V. Bucci, Commensal antimicrobial resistance mediates microbiome resilience to antibiotic disruption. Sci. Transl. Med. 16, eadi9711 (2024). doi: 10.1126/scitranslmed.adi9711.

19. M. Bakhit, T. Hoffmann, A.M. Scott, E. Beller, J. Rathbone, C. Del Mar, Resistance decay in individuals after antibiotic exposure in primary care: a systematic review and meta-analysis. BMC Medicine 16, 126 (2018). doi: 10.1186/s12916-018-1109-4.

20. C. Roubaud-Baudron, V.E. Ruiz, A.M. Swan, B.A. Vallance, C. Ozkul, Z. Pei, J. Li, T.W. Battaglia, G.I. Perez-Perez, M.J. Blaser, Long-term effects of early-life antibiotic exposure on resistance to subsequent bacterial infection. mBio 10, e02820–19 (2019). doi: 10.1128/mBio.02820-19.

21. C. Liu, E.L. Cyphert, S.J. Stephen, B. Wang, A.L. Morales, J.C. Nixon, N.R. Natsoulas, M. Garcia, P. Blazquez Carmona, A.C. Vill, E. Donnelly, I.L. Brito, D. Vashishth, C.J. Hernandez, Microbiome-induced increases and decreases in bone matrix strength can be initiated after skeletal maturity. J. Bone Miner. Res. 39, 1621–1632 (2024). doi: 10.1093/jbmr/zjae157.

22. W.E. Anthony, B. Wang, K.V. Sukhum, A.W. D’Souza, T. Hink, C. Cass, S. Seiler, K.A. Reske, C. Coon, E.R. Dubberke, C.A.D. Burnham, G. Dantas, J.H. Kwon, Acute and persistent effects of commonly used antibiotics on the gut microbiome and resistome in healthy adults. Cell Rep. 39, 110649 (2022). doi: 10.1016/j.celrep.2022.110649.

23. J de la Cuesta-Zuluaga, S.T. Kelley, Y. Chen, J.S. Escobar, N.T. Mueller, R.E. Ley, D. McDonald, S. Huang, A.D. Swafford, R. Knight, V.G. Thackray, Age- and sex-dependent patterns of gut microbial diversity in human adults. mSystems 4, e00261–19 (2019). doi: 10.1128/mSystems.00261-19.

24. L.E. Bryan, S.K. Kowand, H.M. Van Den Elzen, Mechanism of aminoglycoside antibiotic resistance in anaerobic bacteria: Clostridium perfringens and Bacteroides fragilis. Antimicrob. Agents Chemother. 15, 7–13 (1979). doi: 10.1128/aac.15.1.7.

25. R. Edwards, Resistance to beta-lactam antibiotics in Bacteroides spp. J. Med. Microbiol. 46, 979–986 (1997). doi: 10.1099/00222615-46-12-979.

26. D.J. Schwartz, A.E. Langdon, G. Dantas, Understanding the impact of antibiotic perturbation on the human microbiome. Genome Med. 12, 82 (2020). doi: 10.1186/s13073-020-00782-x.

27. M. Derrien, A.S. Alvarez, W.M. de Vos, The gut microbiota in the first decade of life. Trends Microbiol. 27, 997–1010 (2019). doi: 10.1016/j.tim.2019.08.001.

28. A. Parker, S. Romano, R. Ansorge, A. Aboelnour, G. Le Gall, G.M. Savva, M.G. Pontifex, A. Telatin, D. Baker, E. Jones, D. Vauzour, S. Rudder, L.A. Blackshaw, G. Jeffery, S.R. Carding, Fecal microbiota transfer between young and aged mice reverses hallmarks of the aging gut, eye, and brain. Microbiome 10, 68 (2022). doi: 10.1186/s40168-022-01243-w.

29. D.V. Patangia, G. Grimaud, C.A. O’Shea, C.A. Ryan, E. Dempsey, C. Stanton, R.P. Ross, Early life exposure of infants to benzylpenicillin and gentamicin is associated with a persistent amplification of the gut resistome. Microbiome 12, 19 (2024). doi: 10.1186/s40168-023-01732-6.

30. T. Tavella, S. Turroni, P. Brigidi, M. Candela, S. Rampelli, The human gut resistome up to extreme longevity. mSphere 6, e00691–21 (2021). doi: 10.1128/msphere.00691-21.

31. H. Todman, S. Arya, M. Baker, D.J. Stekel, A model of antibiotic resistance genes accumulation through lifetime exposure from food intake and antibiotic treatment. PLoS One 18, e0289941 (2023). doi: 10.1371/journal.pone.0289941.

32. D. Baur, B.P. Gladstone, F. Burkert, E. Carrara, F. Foschi, S. Döbele, E. Tacconelli, Effect of antibiotic stewardship on the incidence of infection and colonization with antibiotic-resistant bacteria and Clostridium difficile infection: a systematic review and meta-analysis. Lancet Infect. Dis. 17, 990–1001 (2017). doi: 10.1016/S1473-3099(17)30325-0.

33. C. Munck, R.U. Sheth, E. Cuaresma, J. Weidler, S.L. Stump, P. Zachariah, D.H. Chong, A.C. Uhlemann, J.A. Abrams, H.H. Wang, D.E. Freedberg, The effect of short-course antibiotics on the resistance profile of colonizing gut bacteria in the ICU: a prospective cohort study. Crit. Care 24, 404 (2020). doi: 10.1186/s13054-020-03061-8.

34. Y. Hong, H. Li, L. Chen, H. Su, B. Zhang, Y. Luo, C. Li, Z. Zhao, Y. Shao, L. Guo, Short-term exposure to antibiotics begets long-term disturbance in gut microbial metabolism and molecular ecological networks. Microbiome 12, 80 (2024). doi: 10.1186/s40168-024-01795-z.

35. C.W. MacPherson, O. Mathieu, J. Tremblay, J. Champagne, A. Nantel, S.A. Girard, T.A. Tompkins, Gut bacterial microbiota and its resistome rapidly recover to basal state levels after short-term amoxicillin-clavulanic acid treatment in healthy adults. Sci. Rep. 8, 11192 (2018). doi: 10.1038/s41598-018-29229-5.

36. B.G. Bell, F. Schellevis, E. Stobberingh, H. Goossens, M. Pringle, A systematic review and meta-analysis of the effects of antibiotic consumption on antibiotic resistance. BMC Infect. Dis. 14, 13 (2014). doi: 10.1186/1471-2334-14-13.

37. K. Kandelaki, C.S. Lundborg, G. Marrone, Antibiotic use and resistance: a cross-sectional study exploring knowledge and attitudes among school and institution personnel in Tbilisi, Republic of Georgia. BMC Res. Notes 8, 495 (2015). doi: 10.1186/s13104-015-1477-1.

38. S.W. Olesen, M.L. Barnett, D.R. MacFadden, J.S. Brownstein, S. Hernández-Díaz, M. Lipsitch, Y.H. Grad, The distribution of antibiotic use and its association with antibiotic resistance. eLife 7, e39435 (2018). doi: 10.7554/eLife.39435.

39. A. Dhariwal, L.C.H. Braten, K. Sturød, G. Salvadori, A. Bargheet, H. Amdal, R. Junges, D. Berild, J.A. Zwart, K. Storheim, F.C. Petersen, Differential response to prolonged amoxicillin treatment: long-term resilience of the microbiome versus long-lasting perturbations in the gut resistome. Gut Microbes 15, 2157200 (2023). doi: 10.1080/19490976.2022.2157200.

40. K. Kang, L. Imamovic, M.A. Misiakou, M.B. Sørensen, Y. Heshiki, Y. Ni, T. Zheng, J. Li, M.M.H. Ellabaan, M. Colomer-Lluch, A.A. Rode, P. Bytzer, G. Panagiotou, M.O.A. Sommer, Expansion and persistence of antibiotic-specific resistance genes following antibiotic treatment. Gut Microbes 13, 1–19 (2021). doi: 10.1080/19490976.2021.1900995.

41. C. d’Humieres, M. Delavy, L. Alla, F. Ichou, E. Gauliard, A. Ghozlane, F. Levenez, N. Galleron, B. Quinquis, N. Pons, J. Mullaert, A. Bridier-Nahmias, B. Condamine, M. Touchon, D. Rainteau, A. Lamaziere, P. Lesnik, M. Ponnaiah, M. Lhomme, N. Sertour, S. Devente, J.D. Docquier, M.E. Bougnoux, O. Tenaillon, M. Magnan, E. Ruppé, N. Grall, X. Duval, D. Ehrlich, F. Mentré, E. Denamur, E.P.C. Rocha, E. Le Chatelier, C. Burdet for the PrediRes study group, Perturbation and resilience of the gut microbiome up to 3 months after β-lactams exposure in healthy volunteers suggest an important role of microbial β-lactamases. Microbiome 12, 50 (2024). doi: 10.1186/s40168-023-01746-0.

42. L. Liu, M.E. Kirst, L. Zhao, E. Li, G.P. Wang, Microbiome resilience despite a profound loss of minority microbiota following clindamycin challenge in humanized gnotobiotic mice. Microbiol. Spectr. 10, e0196021 (2022). doi: 10.1128/spectrum.01960-21.

43. Y. Hu, X. Yang, N. Lu, B. Zhu, The abundance of antibiotic resistance genes in human guts has correlation to the consumption of antibiotics in animal. Gut Microbes 5, 245–249 (2014). doi: 10.4161/gmic.27916.

44. J.R. Huddleston, Horizontal gene transfer in the human gastrointestinal tract: potential spread of antibiotic resistance genes. Infect. Drug Resist. 7, 167–176 (2014). doi: 10.2147/IDR.S48820.

45. K. Forslund, S. Sunagawa, J.R. Kultima, D.R. Mende, M. Arumugam, A. Typas, P. Bork, Country-specific antibiotic use practices impact the human gut resistome. Genome Res. 23, 1163–1169 (2013). doi: 10.1101/gr.155465.113.

46. M.H. Woodworth, R.E. Conrad, M. Haldopoulos, S.M. Pouch, A. Babiker, A.K. Mehta, K.L. Sitchenko, C.H. Wang, A. Strudwick, J.M. Ingersoll, C. Philippe, S. Lohsen, K. Kocaman, B.G. Lindner, J.K. Hatt, R.M. Jones, C. Miller, A.S. Neish, R. Friedman-Moraco, G. Karadkhele, K.H. Liu, D.P Jones, C.C. Mehta, T.R. Ziegler, D.S. Weiss, C.P. Larsen, K.T. Konstantinidis, C.S. Kraft, Fecal microbiota transplantation promotes reduction of antimicrobial resistance by strain replacement. Sci. Transl. Med. 15, eabo2750 (2023). doi: 10.1126/scitranslmed.abo2750.

47. J. Hyun, S.K. Lee, J.H. Cheon, D.E. Yong, H. Koh, Y.K. Kang, M.H. Kim, Y. Sohn, Y. Cho, Y.J. Baek, J.H. Kim, J.Y. Ahn, S.J. Jeong, J.S. Yeom, J.Y. Choi, Faecal microbiota transplantation reduces amounts of antibiotic resistance genes in patients with multidrug-resistant organisms. Antimicrob. Resist. Infect. Control 11, 20 (2022). doi: 10.1186/s13756-022-01064-4.

48. K.L. Kalantar, T. Carvalho, C.F.A. de Bourcy, B. Dimitrov, G. Dingle, R. Egger, J. Han, O.B. Holmes, Y.F. Juan, R. King, A. Kislyuk, M.F. Lin, M. Mariano, T. Morse, L.V. Reynoso, D.R. Cruz, J. Sheu, J. Tang, J. Wang, M.A. Zhang, E. Zhong, V. Ahyong, S. Lay, S. Chea, J.A. Bohl, J.E. Manning, C.M. Tato, J.L. DeRisi, IDseq-an open source cloud-based pipeline and analysis service for metagenomic pathogen detection and monitoring. Gigascience 9, giaa111 (2020). doi: 10.1093/gigascience/giaa111.

49. B.W. Ma, N.A. Bokulich, P.A. Castillo, A. Kananurak, M.A. Underwood, D.A. Mills, C.L. Bevins, Routine habitat change: a source of unrecognized transient alteration of intestinal microbiota in laboratory mice. PLoS ONE 7, e47416 (2012). doi: 10.1371/journal.pone.0047416.

50. K.E. Fujimura, A.R. Sitarik, S. Havstad, D.L. Lin, S. Levan, D. Fadrosh, A.R. Panzer, B. LaMere, E. Rackaityte, N.W. Lukacs, G. Wegienka, H.A. Boushey, D.R. Ownby, E.M. Zoratti, A.M. Levin, C.C. Johnson, S.V. Lynch, Neonatal gut microbiota associates with childhood multisensitized atopy and T cell differentiation. Nat. Med. 22, 1187–1191 (2016). doi: 10.1038/nm.4176.

